# Critical role of PCYT2 in muscle health and aging

**DOI:** 10.1101/2022.03.02.482658

**Authors:** Domagoj Cikes, Kareem Elsayad, Erdinc Sezgin, Erika Koitai, Torma Ferenc, Michael Orthofer, Rebecca Yarwood, Leonhard X. Heinz, Vitaly Sedlyarov, Nasser Darwish Miranda, Adrian Taylor, Sophie Grapentine, Fathiya al-Murshedi, Anne Abott, Adelheid Weidinger, Candice Kutchukian, Colline Sanchez, Shane J.F. Cronin, Maria Novatchkova, Anoop Kavirayani, Thomas Schuetz, Bernhard Haubner, Lisa Haas, Astrid Hagelkruys, Suzanne Jackowski, Andrey Kozlov, Vincent Jacquemond, Claude Knauf, Giulio Superti-Furga, Eric Rullman, Thomas Gustafsson, John McDermot, Martin Lowe, Zsolt Radak, Jeffrey S. Chamberlain, Marica Bakovic, Siddharth Banka, Josef M. Penninger

## Abstract

Muscle degeneration is the most prevalent cause for frailty and dependency in inherited diseases and ageing, affecting hundreds of millions of people. Elucidation of pathophysiological mechanisms, as well as effective treatments for muscle diseases represents an important goal in improving human health. Here, we show that phosphatidylethanolamine cytidyltransferase (PCYT2/ECT), the critical enzyme of the Kennedy branch of phosphatidylethanolamine (PE) synthesis pathway, has an essential role in muscle health. Human genetic deficiency in *PCYT2* causes a severe disease with failure to thrive and progressive muscle weakness. *Pcyt2* mutant zebrafish recapitulate the patient phenotypes, indicating that the role of PCYT2/PE in muscle is evolutionary conserved. Muscle specific *Pcyt2* knockout mice exhibited failure to thrive, impaired muscle development, progressive muscle weakness, muscle loss, accelerated ageing, and reduced lifespan. Mechanistically, Pcyt2 deficiency affects mitochondrial bioenergetics and physicochemical properties of the myofiber membrane lipid bilayer, in particular under exercise strain. We also show that PCYT2 activity declines in the aging muscles of humans and mice. AAV-based delivery of PCYT2 rescued muscle weakness in *Pcyt2* knock-out mice and, importantly, improved muscle strength in old mice, offering a novel therapeutic avenue for rare disease patients and muscle aging. Thus, PCYT2 plays a fundamental, specific, and conserved role in vertebrate muscle health, linking PCYT2 and PCYT2 synthesized PE lipids to severe muscle dystrophy, exercise intolerance and aging.

## Summary

Skeletal muscle is the biggest organ in the human body, with essential roles in mechanical support, mobility and energy expenditure. Loss and degeneration of muscle tissue, either as a result of inherited diseases ^1^, chronic diseases, or ageing ^2^ severely impairs the life quality, independence, and health, of millions of people world-wide. Therefore, complete understanding of pathophysiological mechanisms that affect muscle tissue, as well as development of novel treatments, represents an important objective in medicine.

The eukaryotic lipidome is uniquely complex with a potential of generating up to 100 000 different lipid species ^3^. Differences among individual lipid species occur at subcellular compartments and between cell and tissue types ^4^. Tissue dependent differences in lipid composition suggest that certain organs use specific lipid synthesis pathways, essential for organ health and longevity ^5^.

In humans, genetic deficiency in phosphatidylethanolamine cytidyltransferase (PCYT2/ECT), the bottle neck enzyme in synthesis of PE through the Kennedy pathway ^6^, leads to complex and severe hereditary spastic paraplegia (HSP) ^7,8^. Here, we discover a conserved, essential and specific role for PCYT2 synthesized phosphatidylethanolamine (PE) in muscle health. *Pcyt2* mutant Zebrafish and muscle specific *Pcyt2* knockout mice recapitulate several patient phenotypes, particularly failure to thrive, short stature, impaired muscle development, progressive weakness, inflammation, and accelerated ageing, resulting in shortened life spans. In contrast, mice lacking Pcyt2 in other tissues appeared unaffected. Loss of PCYT2 in muscle results in alterations in the mitochondrial and cellular lipidome, affecting mitochondrial activity and physicochemical properties of the lipid bilayer, compromising sarcolemmal stability that is further aggravated by a mechanical strain. We further show that PCYT2 activity declines in aging muscles of humans and mice and that *Pcyt2* gene-therapy in *Pcyt2* knock-out mice and in aged mice improved muscle strength. Thus, PCYT2 and PE, synthesized via Pcyt2, play fundamental roles in muscle biology, linking mitochondrial homeostasis and sarcolemmal lipid bilayer disorders to muscle degeneration, exercise strain tolerance and aging.

### Patients with disease-causing *PCYT2* variants fail to thrive

*PCYT2* mutations were recently discovered in human patients who manifest a complex disorder that involves developmental gross motor delay and progressive overall muscle weakness ^7^. Observing these patients, we found that those with a homozygous nonsense variant NM_001184917.2:c.1129C>T (p.Arg377Ter) in *PCYT2* exhibited a significantly lower weight and shorter body length starting from birth and continuing throughout childhood, puberty and early adulthood (Figure 1A,B). We also assessed patients with mutations in *EPT1*, which encodes the enzyme catalyzing the next and final step in PE synthesis via the Kennedy pathway (Extended Figure 1A,B). These patients have been reported to manifest similar clinical features as those with *PCYT2* mutations ^8,9^. Indeed, patients with a homozygous variant NM_033505.4:c.335G>C (p.Arg112Pro) in EPT1 also exhibited growth defects compared to unaffected children, further indicating a role for the Kennedy pathway in growth and development (Extended Data Figure 1B). Thus, in addition to previously described symptoms such as progressive neurological deterioration, mutations in two critical enzymes that generate PE in the Kennedy pathway are associated with stunted growth starting from birth and continuing throughout childhood.

**Figure 1.**
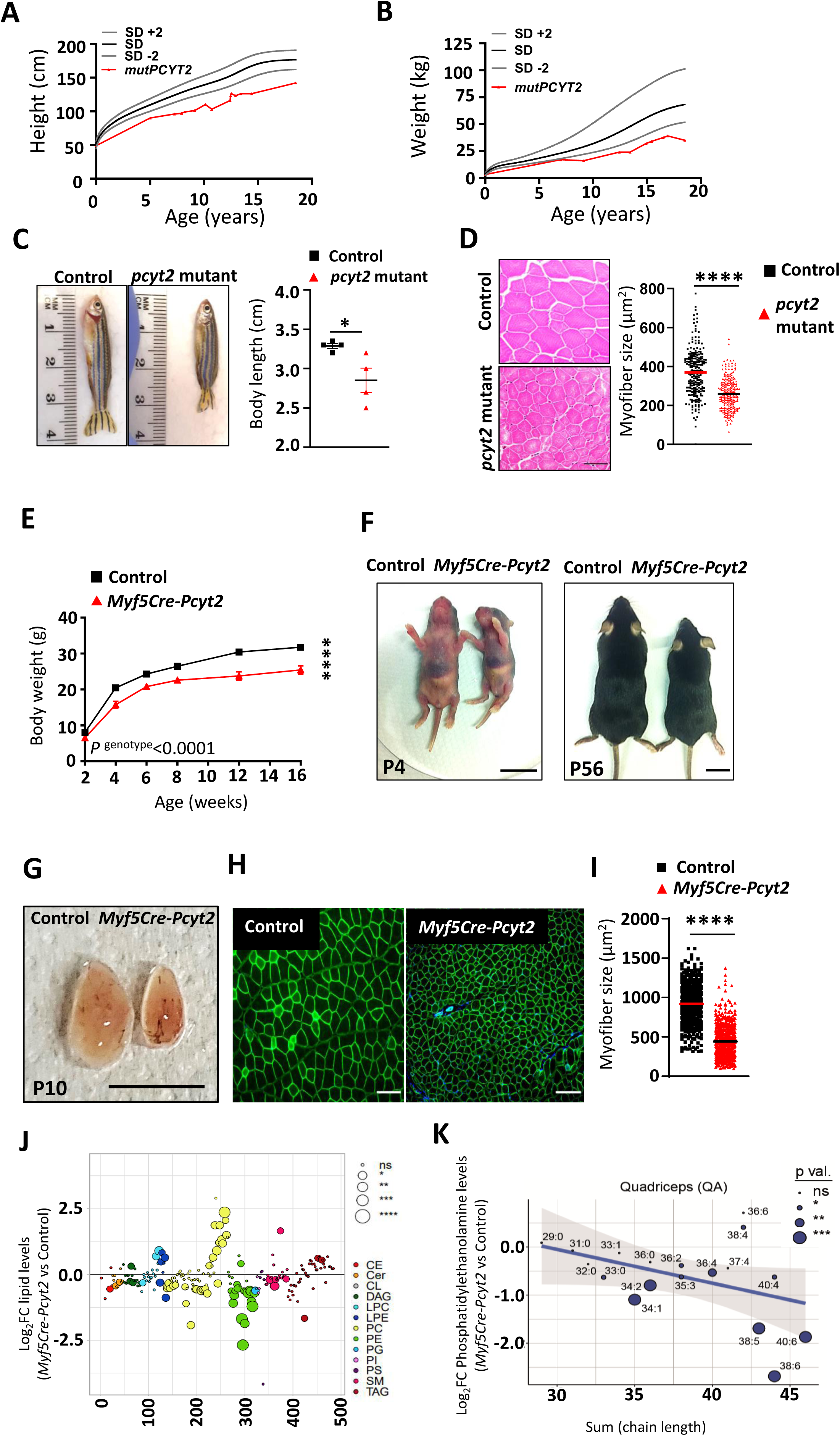
Phenotypes of human *PCYT2* rare disease mutation and *pcyt2* mutant zebrafish. **(A)** Body weight and **(B)** height gain of patient carrying the homozygous nonsense variant 3c.1129C>T (p.Arg377Ter) in the *PCYT2* gene. Controls indicate WHO standards of median weights and heights at the respective ages +/-2 standard deviations (SD). **(C)** Representative appearance and quantifications of body length of control and hypomorphic *pcyt2* mutant zebrafish at 14 months post fertilization. n=4 for each group. **(D)** Representative muscle sections and muscle myofiber sizes of control and hypomorphic pcyt2 zebrafish. Scale bar 50µm. N≥ 70 myofibers of the same anatomical region were analyzed, n=4 animals per group **(E)** Body weight gains of control and *Myf5Cre-Pcyt2* littermates on standard chow diet. n=11-15 per group. Two-Way ANOVA with multiple comparison followed by Bonferroni correction was used. ****p^(genotype)^ < 0.0001 **(F)** Appearance of 4 day old (P4) and 56 day old (P56) control and *Myf5Cre-Pcyt2* littermates. Scale bars are 1 cm for P4 and 2 cm for P56. **(G)** Skeletal muscle appearance (quadriceps) isolated from 10 day control and *Myf5Cre-Pcyt2* littermate mice. (H) Representative cross sections and distribution of skeletal muscle myofiber diameter sizes from 6 month old control and *Myf5Cre-Pcyt2* mice. Myofibers were imaged using 10X magnification with ≥ 100myofibers analyzed per mouse. n=4 animals per group. Scale bar 100µm. **(I)** Lipidomics analyses from quadriceps muscles isolated from 10 day old *Myf5Cre-Pcyt2* and littermate control mice. Data are shown relative to control values. CE-cholesterol ester; Cer-Ceramides; DAG-diacylglycerols; LPC-lysophosphatidylcholines; LPE-lysophosphatidylethanolamines; PC-phosphatidylcholines; PE-phosphatidylethanolamines; PG-phosphatidylglycerols; PI-phosphatidylinositols; PS-phosphatidylserines; SM-sphingomyelins; TAG-triacylglycerols. n=4 per group. **(J)** Detailed analysis of PE species with different chain lengths from quadriceps muscles of *Myf5Cre-Pcyt2* as compared to control mice.

Data are shown as means ± SEM. *p < 0.05, **p < 0.01, ***p < 0.001, and ****p < 0.0001, n.s. not significant. Unpaired Student t-test with Welch correction was used for statistical analysis unless stated otherwise.

### *Pcyt2* deficiency in zebrafish results in stunted growth and small myofibers

Since Ept1 loss can be partially compensated by Cept1 ^10^, we therefore focused on the bottleneck enzyme PCYT2 (Extended Figure 1A). Given the ubiquitous tissue expression of PCYT2 ^11^, its loss of function could potentially affect several organs, thus contributing to the complexity and severity of the disease. To gain insight into pathophysiological mechanisms, we first examined hypomorphic mutant *pcyt2* zebrafish generated using the CRISPR/Cas9 system ^7^. Similar to the human rare disease patients, *pcyt2* mutant zebrafish were significantly smaller than wild type (Figure 1C).

Zebrafish and mouse models of hereditary spastic paraplegia rarely exhibit whole body growth phenotypes ^12,13^ ^14,15^. In contrast, muscle development is essential for whole-body growth, and failure to thrive with short stature are well known features of muscular dystrophies ^16^. Therefore, we examined muscle morphology in *pcyt2* mutant zebrafish. We observed significantly smaller skeletal muscle fibers and reduced myofiber diameters in *pcyt2* mutant zebrafish compared to controls (Figure 1D). These results suggest that reduced muscle size could explain the stunted growth associated with *pcyt2* loss-of-function mutations in zebrafish and patients.

### *Pcyt2* muscle deficiency in mice impairs early muscle growth and development

In mice, global disruption of Pcyt2 results in embryonic lethality ^17^. Therefore, to study the role of Pcyt2 in muscle physiology and development, we generated mice with a muscle-specific *Pcyt2* deletion during early muscle development to recapitulate the human condition. Briefly, we crossed *Pcyt2^flox/flox^* with *Myf5* promoter driven *Cre* mice to generate *Myf5Cre-Pcyt2* offspring, given that Myf5 is the first broadly expressed myogenic regulatory factor in the developing myotome ^18^. *Pcyt2* deletion was validated by RNA sequencing (Extended Data Figure 2A, *Myf5Cre-Pcyt2* mice were born at normal Mendelian ratios, but were significantly smaller at birth (postnatal day 1, P1) and early postnatal days (P4), gained less weight, and grew significantly less in length during the postnatal period compared to controls, as observed for both genders (Figure 1E,F; Extended Data Figure 3A-D). Neither *Myf5Cre* nor *Pcyt2^flox/flox^*littermate controls displayed a phenotype, therefore we used *Pcyt2^flox/flox^*littermates as controls for all subsequent experiments. We noticed that limb muscles were smaller in *Myf5Cre-Pcyt2* mice compared to controls already at P10 and still at 2 months old (Figure 1G; Extended Data Figure 3E-H). Myofiber size was significantly reduced in skeletal muscle (Figure 1G,I). Lipidomic analysis of quadriceps muscle isolated from 10-day old *Myf5Cre-Pcyt2* mice showed a marked and specific reduction in the levels of PE, particularly of long-chain fatty acid PE species (Figure 1J, K).

**Figure 2.**
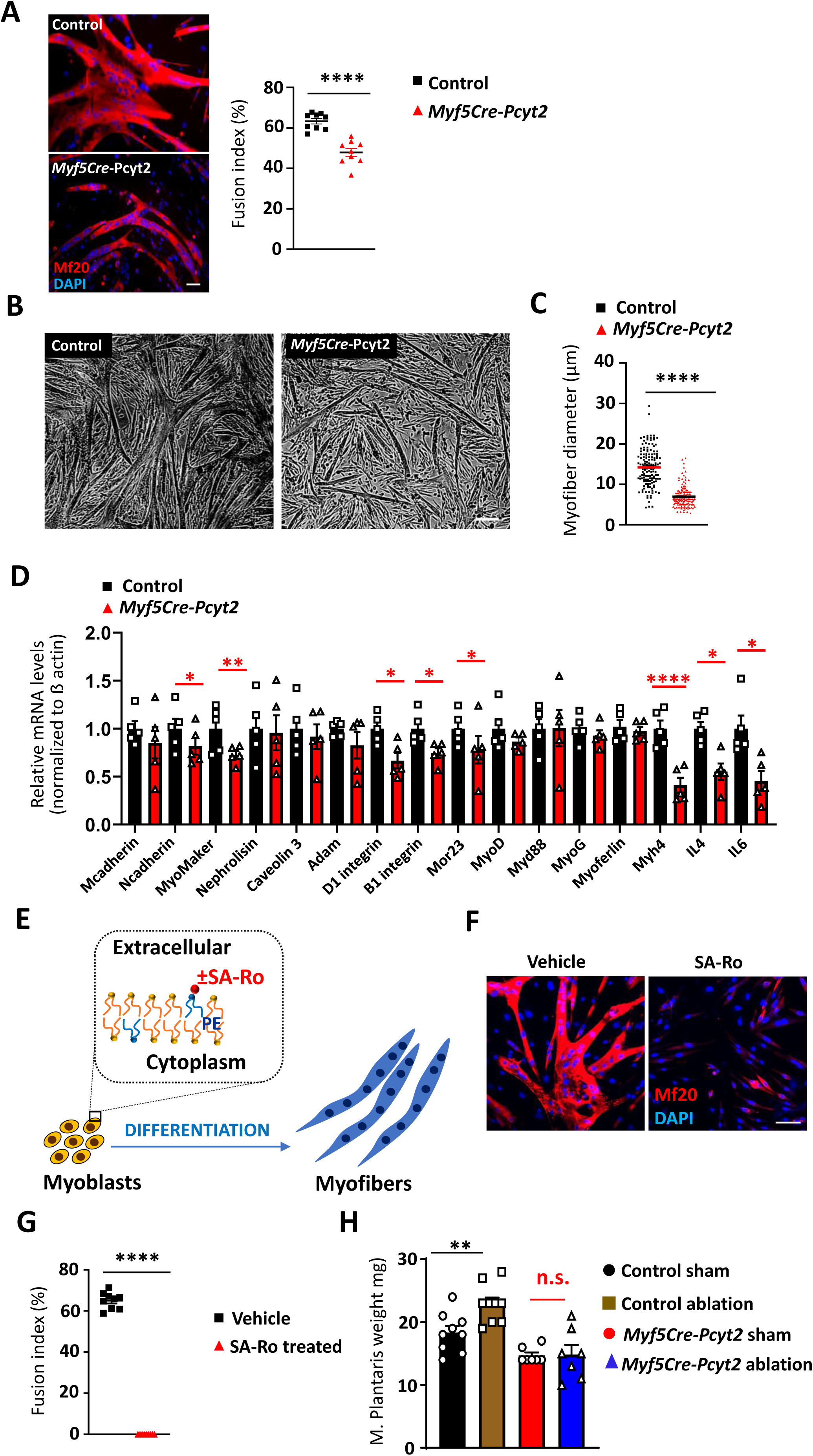
*Pcyt2* deficiency affects muscle stem cell fusion and muscle hypertrophic growth. **(A)** Representative images and **(B)** myofiber diameter quantification of Control and *Myf5Cre-Pcyt2* myoblasts after differentiation *in vitro*. Myofibers were imaged in triplicates using 10X magnification. Each dot represents myofiber diameter with a total of ≥ 50 counted myofibers per one cell culture. N=3 animals per group. Scale bar 50µm. **(C)** Representative images of Mf20 stained myofibers and primary myoblast fusion index quantification of Control and *Myf5Cre-Pcyt2* primary myoblasts after differentiation *in vitro*. Each dot represents a calculated fusion index (number of fused nuclei/total number of nuclei) in triplicates, with a total of ≥ 300 counted nuclei per one isolation. Myofibers were imaged using 10X magnification. N=3 animals per group. Scale bar 50µm. **(D)** RT-PCR analysis of fusion and differentiation markers of Control and Myf5Cre-Pcyt2 myoblasts after 48h in differentiation media. N=5 cell cultures from 5 different animals per group. **(E)** Representative images and myoblast fusion index quantification of primary myoblasts with addition of vehicle (DMSO) and SA-Ro phosphatidylethanolamine binding peptide in differentiation media. Each dot represents a calculated fusion index (number of fused nuclei/total number of nuclei) in triplicates, with a total of ≥ 300 counted nuclei per one isolation. Myofibers were imaged using 10X magnification. Scale bar 50µm. **(F)** Hypertrophic muscle growth in control and Myf5Cre-Pcyt2 mice. Following synergic ablation or sham surgery, M. plantaris weights were determined on the compensating limb. Each dot represents individual mice. Data are shown as means ± SEM. *p < 0.05, **p < 0.01, ***p < 0.001, and ****p < 0.0001, n.s. not significant. Unpaired Student t-test with Welch correction was used for statistical analysis unless stated otherwise.

**Figure 3.**
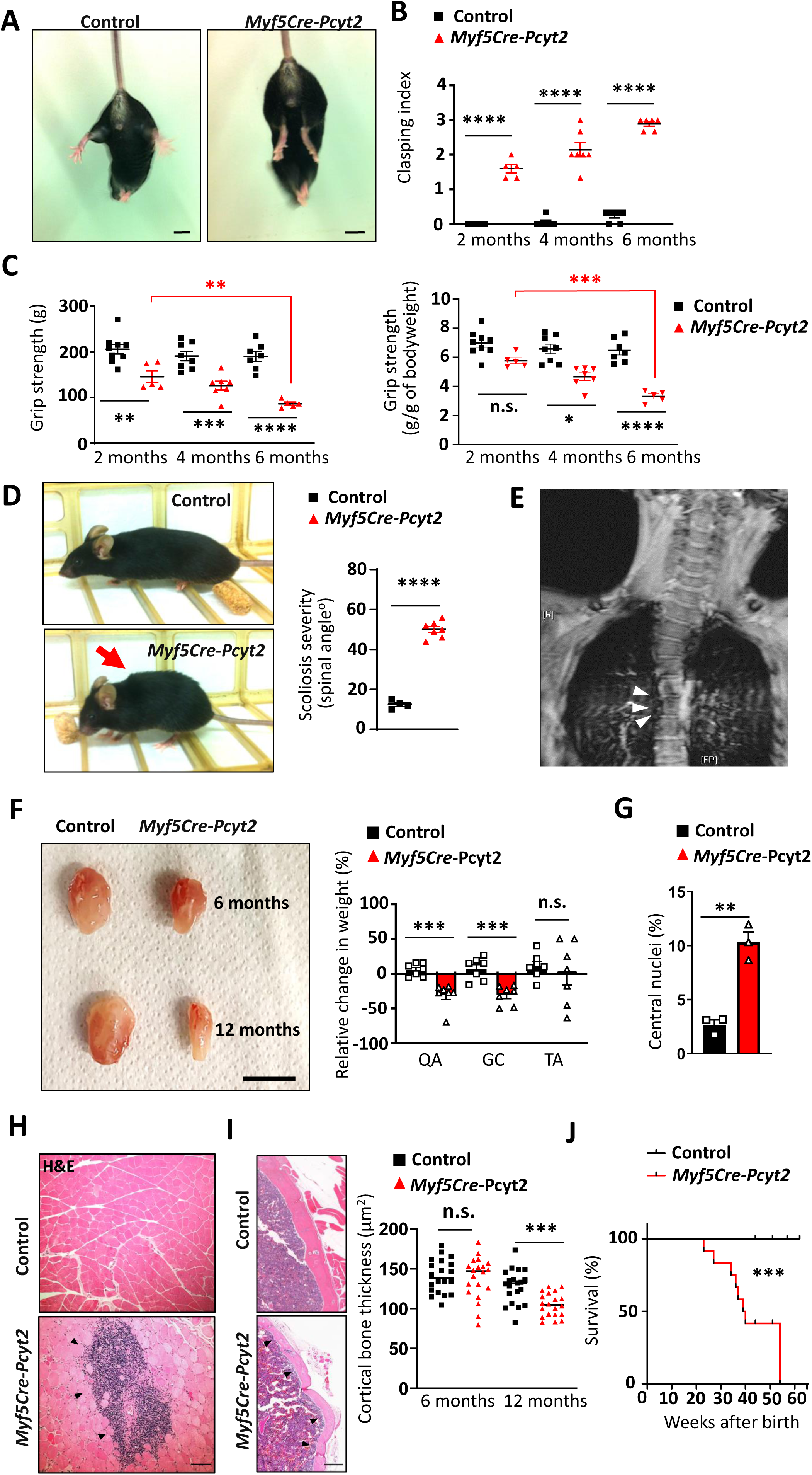
Inactivation of *Pcyt2* in mice leads to progressive weakness, muscle atrophy, inflammation and accelerated ageing. **(A)** Representative images of 6 month old control and *Myf5Cre-Pcyt2* mice and (B) quantification of progressive worsening of hind limb clasping **(B)**. Each dot represents one mouse, values are average of three measurements per mouse; scale bar 1 cm. **(C)** Age-dependent decline in grip strength in control and *Myf5Cre-Pcyt2* littermates. Each dot represents one mouse, values are average of three measurements per mouse. **(D)** Typical kyphosis appearance and kyphosis severity in 8 month old control and *Myf5Cre-Pcyt2* mice. n=4-7 per group. **(E)** Evident scoliosis (arrows) in a patient carrying the homozygous nonsense variant NM_001184917.2:3c.1129C>T (p.Arg377Ter) in *PCYT2*. **(F)** Representative image and quantification of relative muscle mass changes of 12 month old versus 6 month old control and *Myf5Cre-Pcyt2* littermates. QA, quadriceps; GC, gastrocnemius; TA, tibialis anterior muscles. Scale bar = 1 cm; n=7 per group. **(G)** Quantification of myofibers with central nuclei in quadriceps muscles from 8 month old control and *Myf5Cre-Pcyt2* mice. n=3 mice per group. Scale bar 100µm. **(H)** Muscle inflammation as determined by H&E staining. Data are from 12 month old mice. Data are representative for n=4 mice per group. Scale bar 100μm. **(I)** Representative cross section of tibial bone in 12 month old control and *Myf5Cre-Pcyt2* mice with quantification of tibial bone cortical thickness. Randomly assigned 5 areas from n=4 mice per group were quantified. **(J)** Survival curves for control and *Myf5Cre-Pcyt2* mice. n=22 mice per group. For statistical analysis Mantel Cox test). Data are shown as means ± SEM. Each dot represents individual mice. *p < 0.05, **p < 0.01, ***p < 0.001, and ****p < 0.0001, n.s. not significant Unpaired Student t-test with Welch correction was used for statistical analysis unless stated otherwise.

Muscle growth is mediated first by muscle satellite cell proliferation and an increase in myofibers until ∼P7, and subsequently via myofiber hypertrophy ^19^. The number of proliferating cells as well as the rate of myoblast proliferation was not affected, as inferred from BrdU incorporation in the developing muscles of *Myf5Cre-Pcyt2* mice and EdU incorporation in proliferating myoblasts *in vitro* (Extended Data Figure 4A, B). Although the number and distribution of Pax7^+^ muscle progenitor cells was similar in adult *Myf5Cre-Pcyt2* and control mice (Extended Data Figure 4C), *Myf5Cre-Pcyt2* mice showed a mild but significant reduction in myoblast fusion, with thinner myofibers (Figure 2A-C). Several pro-fusion and differentiation markers were significantly reduced in the myoblasts from *Myf5Cre-Pcyt2* mice (Figure 2D). Of, note There were no apparent changes in the levels of transcriptional regulators of myofiber differentiation, such as MyoD and MyoG (Figure 2D). Membrane lipid composition is essential for myoblast fusion ^20^. Given that PE lipids are abundant membrane lipids, we wanted to address if unavailable PE in the myoblast membrane would affect myoblast fusion into myofibers. We used the non-toxic, PE-specific SA-Ro binding probe ^21^ to mask the externally exposed membrane PE during the process of myoblast fusion, mimicking PE deficiency. Indeed, myoblast fusion was severely affected (Figure 2E), showing that membrane PE is required for efficient myoblast fusion.

**Figure 4.**
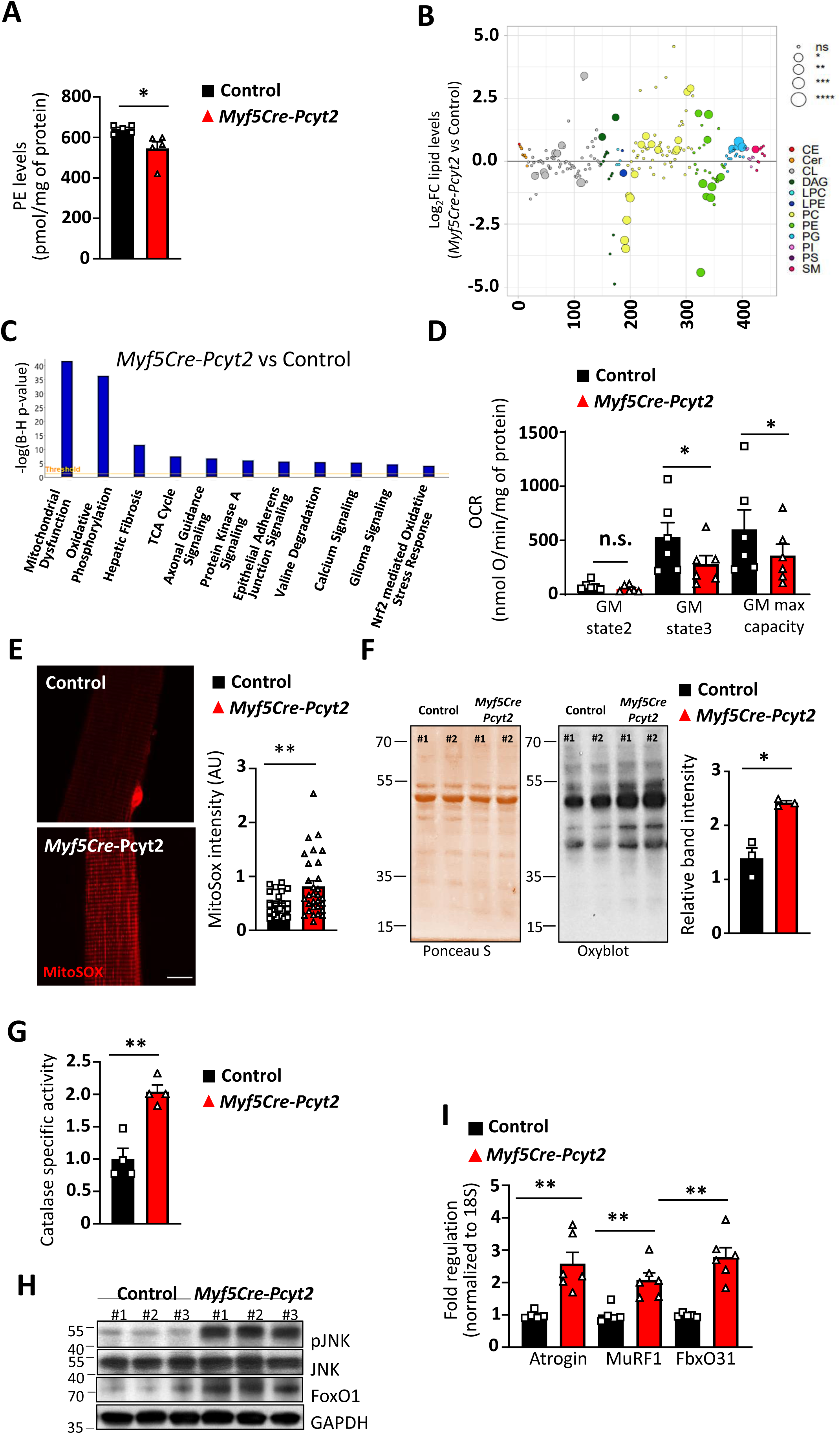
*Pcyt2* deficiency severely affects muscle mitochondrial homeostasis as opposed to brown fat mitochondria. **(A)** Lipidomics analyses of purified mitochondria isolated from 2 month old Control and Myf5Cre-Pcyt2 mice. N=6 mice per group **(B)** Pathway enrichment analysis of differentially expressed genes in Control and Myf5Cre-Pcyt2 quadriceps isolated from 10 day old pups. Evident enrichment of mitochondrial dysfunction linked genes specifically in the muscle of Myf5Cre-Pcyt2 mice. N=4 mice per group. **(C-D)** Muscle mitochondrial function assessed by measurements of complex I linked activity on isolated mitochondria from 2 months and 6 months old control and *Myf5Cre-Pcyt2* mice respectively. Paired Student t-test. (E) Measurement and quantification of myofiber mitochondrial reactive oxygen species (mtROS) in isolated myofibers (EDL muscle) from 6 months old control and *Myf5Cre-Pcyt2* mice. Each dot represents relative amount of mtROS from a single myofiber. N=3 mice per group with ≥10 myofibers analyzed per mouse. **(F)** Evidence of increased protein oxidative damage in quadriceps muscles isolated from 6 months old *Myf5Cre-Pcy*t2 mice. Representative blot are shown for n=3 mice per group. **(H)** Catalase anti-oxidant activity in quadriceps muscles from 6 months old control and *Myf5Cre-Pcyt2* mice. **(I)** Increased levels of phospho-JNK (pJNK) and FoxO1 in quadriceps muscles from 6 months old *Myf5Cre-Pcyt2* mice as compared to controls. n=3 per group. **(J)** Increased levels of myofiber wasting markers in muscles of 8 months old *Myf5Cre-Pcyt2* mice. Each dot represents individual mice. *p < 0.05, **p < 0.01, ***p < 0.001, and ****p < 0.0001, n.s. not significant. Unpaired Student t-test with Welch correction was used for statistical analysis unless stated otherwise.

To directly examine hypertrophic muscle growth, an essential process for muscle growth that is independent of myoblast fusion process, we performed synergic muscle ablation ^22^. Muscle overloading resulted in a significant enlargement of the plantaris muscle on the un-operated limb in control mice but not in *Myf5Cre-Pcyt2* mice (Figure 2F). Phosphorylation of downstream markers of global protein translation and synthesis, ribosomal RPS6 and 4EIF-A, appeared to be unaffected (Extended Data Figure 5A), indicating that altered protein synthesis does not underlie the observed defect in hypertrophic growth of muscles from *Myf5Cre-Pcyt2* mice. The observed impaired myoblast fusion and hypertrophic muscle growth in *Myf5Cre-Pcyt2* mice did not apparently affect the fiber type distribution not fiber number in muscles from adult *Myf5Cre-Pcyt2* mice (Extended Data Figure 5B-D). Together these data show that loss of *Pcyt2* impairs long-chain fatty acid PE production and compromises both progenitor fusion and hypertrophic growth, leading to reduced myofiber sizes. Thus, we infer that *Pcyt2* deficiency in muscle leads to a failure to thrive in zebrafish, mice and humans, indicating a critical evolutionary conserved role for PCYT2 in muscle biology.

**Figure 5.**
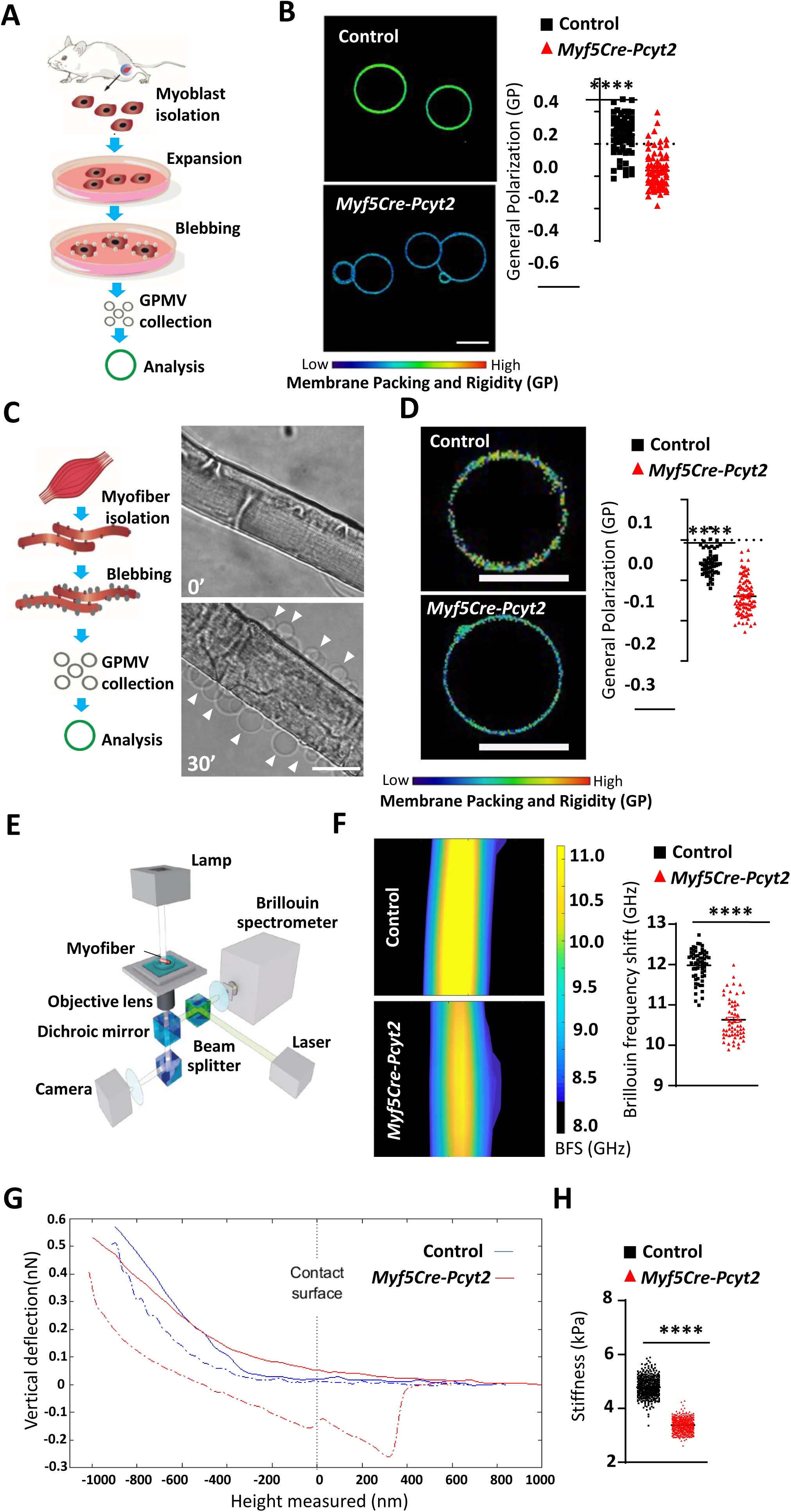
Loss of *Pcyt2* results in altered muscle membrane architectures. **(A)** Schematic diagram of GPMV isolation from primary myoblasts. **(B)** Polarization microscopy analysis of NR12S dye-stained GPMVs isolated from control and *Myf5Cre-Pcyt2* myoblasts. GPMVs were isolated from cultured myoblasts from 2 mice per group. Representative images and quantifications are shown. Each dot represents GP values of a single GPMV. Scale bar 10μm. **(C)** Schematic diagram and representative example of GPMVs (arrows) s immediately after isolation from skeletal myofibers. Images are from a wild type mouse at 0 and 30 minutes under GPMV conditions. Scale bar 50μm. **(D)** Polarization microscopy of NR12S-stained GPMVs isolated from control and *Myf5Cre-Pcyt2* primary myofibers (as shown in **C**). GPMVs were isolated from ≥ 100 myofibers from 2 mice per group. Representative images and quantifications are shown. Each dot represents GP values of a single GPMV. Scale bar 10μm. **(E)** Schematic set-up of Brillouin light scattering microscopy adapted to freshly isolated myofibers. **(F)** Surface stiffness analysis as measured by Brillouin frequency shift (BFS) from isolated control and *Myf5Cre-Pcyt*2 myofibers. Left panels indicate representative microscopy images. Each data point in the right panel represents a BFS peak value of the myofiber surface. > 15myofibers from n=3 mice per group were analyzed. **(G)** Representative qualitative membrane stiffness data of control and *Myf5Cre-Pcyt2* myofibers as assessed by atomic force microscopy. As seen from the curve angles in the approach phase (0 to −1000nm) and retraction phase (−1000 to 0nm), the cantilever bends much less for *Myf5Cre-Pcyt2* myofibers, indicating lower surface stiffness. In the prolonged part of retraction phase (0 to 400nm) the cantilever remains deeper inserted within the *Myf5Cre-Pcyt2* myofibers, indicating a higher degree of surface deformation upon pressure. **(H)** Quantitative myofiber membrane stiffness as assessed by atomic force microscopy to quantify cell membrane stiffness using the Young’s modulus scale (in kilopascal, kPa). For each myofiber we collected ≥4000 measurements over a 5µm X 5µm area. Matlab’s Randsample function was used to uniformly sample each myofiber measurements. Each dot represents 500 data points per each myofiber, from n=20-26 control and *Myf5Cre-Pcyt2* myofibers. Data are shown as means ± SEM. Each dot represents individual mice. *p < 0.05, **p < 0.01, ***p < 0.001, and ****p < 0.0001, n.s. not significant. Unpaired Student t-test with Welch correction was used for statistical analysis unless stated otherwise.

### Muscles lacking *Pcyt2* exhibit progressive muscle wasting

During phenotyping of the mutant mice, we noticed that adult *Myf5Cre-Pcyt2* mice exhibited hindlimb clasping upon tail suspension compared to controls (Figure 3A,B), indicative of muscle weakness. Moreover, muscle strength was significantly reduced and progressively declined as *Myf5Cre-Pcyt2* mice aged (Figure 3C). At 8 months of age all *Myf5Cre-Pcyt2* mice developed kyphosis (Figure 3D), which was also seen in *PCYT2* disease patients (Figure 3E) and has been reported in different mouse models of muscular dystrophy ^23,24^. Severe and progressive loss of muscle tissue was evident in *Myf5Cre-Pcyt2* mice (Figure 2F) with a high incidence of centrally localized nuclei (Figure 3G). Furthermore, we observed tubular aggregates and inflammation in the muscles of 12-15 months old *Myf5Cre-Pcyt2* mice (Figure 3H, Extended Data Figure 6A-C). Consequent to the observed muscle weakness, *Myf5Cre-Pcyt2* mice developed secondary progressive osteopenia contributing to overall frailty (Figure 3I). As a result, *Myf5Cre-Pcyt2* mice had a significantly decreased survival rate compared to controls (Figure 2J).

**Figure 6.**
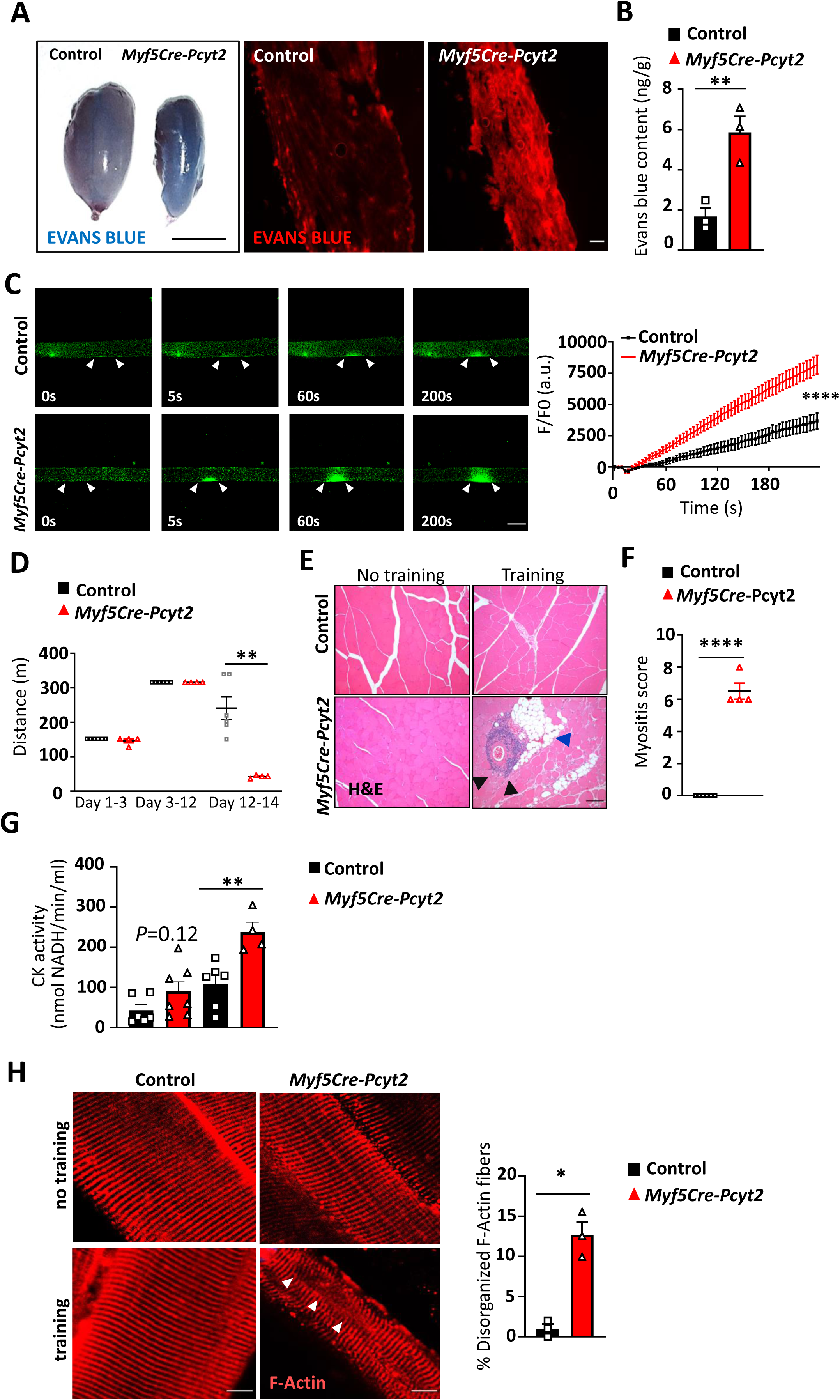
Pcyt2 is essential for muscle membrane integrity and strain tolerance. **(A)** Penetrance of Evans blue into the quadriceps muscle of 6 month old control and *Myf5Cre-Pcyt2* mice after i.p. injection. Gross morphologies left) and histological sections are shown. Scale bars are 1cm and 100µm. **(B)** Quantification of Evans blue in the quadriceps muscle after extraction from the tissue determined n=3 per group. **(C)** Laser induced damage of myofibers isolated from 6 month old control and *Myf5Cre-Pcyt2* mice. The extent of damage was followed over time using influx of the fm43. The injured membrane areas of the myofibers are indicated by two arrows. Right panel shows quantification of fm43 influx over the indicated time period +/-SEM; n=9-12 myofibers per group from two independent experiments. Scale bar 50µm **(D)** Running distance (in meters) during eccentric exercise of 6 month old control and *Myf5Cre-Pcyt2* mice. Data are pooled from the first training bout on days 1-3 (4m min^-1^), the second training bout on days 4-11 (9m min^-1^); and the third training bout on days 12-14 (20m min^-1^). n=4-6 per group. **(E)** Histological analysis (H&E staining) of quadriceps muscles isolated from 6 month old control and *Myf5Cre-Pcyt2 mice*, untrained (no training) and after (training) the eccentric exercise. Black arrows show inflammation migrating from a blood vessel; blue arrow indicates ectopic fat deposits. Scale bars 100µm **(F)** Myopathy scores in 6 month old control and *Myf5Cre-Pcyt2* mice following eccentric exercise. The following parameters were used to calculate the score: inflammatory cell infiltrates, myofiber necrosis, myofiber atrophy, interstitial fibrosis, loss of sarcoplasmic membrane integrity, regenerating myofibers. Each was scored with 1-4 depending of the severity, and summed for the final value, n=4-6 mice per cohort. **(G)** Blood muscle creatine kinase levels as addressed by measurements of muscle creatine kinase activity in sera from 6 month old sedentary and immediately after eccentric exercise regime of control and Myf5Cre-Pcyt2 mice. **(H)** F-actin staining of skeletal muscle tissue isolated from 6 months old control and *Myf5Cre-Pcyt2* mice after eccentric exercise. Images of quadriceps cross-sections were taken using 20x magnifications. ≥100 myofibers were counted. n=3 mice per group. Scale bar 15µm. Each dot represents individual mice. *p < 0.05, **p < 0.01, ***p < 0.001, and ****p < 0.0001, n.s. not significant. Unpaired Student t-test with Welch correction was used for statistical analysis unless stated otherwise.

As expected, myopathic changes and consequent progressive sarcopenia had a marked effect on whole-body metabolism. At 8 months of age, blood glucose levels were significantly lower in *Myf5Cre-Pcyt2* mice; food consumption was also markedly decreased (Extended Data Figure 6D-E). Although *Myf5Cre-Pcyt2* mice were less active (Extended Data Figure 6F), energy expenditure relative to body weight was significantly increased in both light and dark periods (Extended Data Figure 6G). Under thermoneutrality, during the light phase energy expenditure of Myf5Cre Pcyt2 mice was comparable to Control mice, while during the dark phase the higher energy expenditure of Myf5Cre Pcyt2 remained (Extended Data Figure 6H). However, the lower activity and the muscle weakness (grip strength) of *Myf5Cre-Pcyt2* mice were also evident under thermoneutrality conditions (Extended Data Figure 6I,J). These data show that muscles specific *Pcyt2* mutant mice exhibit progressive weakness and skeletal muscle inflammation.

### Pcyt2 is specifically required in muscle

*Myf5Cre* is active in precursors of both skeletal muscle and brown adipose tissue (BAT) (but not heart muscle) ^25^. PE species were also reduced in the BAT of *Myf5Cre-Pcyt2* mice, but to a markedly lower extent than in the skeletal muscle (Extended Data Figure 7A,B). Importantly, loss of *Pcyt2* did not affect *in vitro* differentiation of adipocyte progenitors into brown fat, thermoregulation through BAT activity in cold exposure and fasting, nor levels of the brown fat marker UCP1 in BAT (Extended Data Figure 7C-F). In addition, mitochondrial ultra-structures, content and respiration appeared normal in BAT of *Myf5Cre-Pcyt2* mice (Extended Data Figure 7G-J). Moreover, we crossed *Pcyt2*^flox/flox^ mice to the *AdipoQCre* line to remove *Pcyt2* specifically in white and brown adipose tissue ^26^ and did not observe any differences in growth or blood glucose, nor any apparent pathologies (Extended Data Figure 8A-C). Tissue-specific deletion of *Pcyt2* in motor neurons (*Mnx1Cre*; Extended Data Figure 8D-F), gut epithelium (*Villin1Cre*; Extended Data Figure 8G-I), and epithelial cells within the mammary gland and skin (*K14Cre*, Extended Data Figure 8J-L), neither resulted in any apparent developmental deficiencies nor any degenerative phenotypes up to 12 months of age. This lack of any apparent phenotypes in the above tested Cre-deleter mouse lines suggests that Pcyt2 does not play an essential role in the plasma or mitochondrial membranes of these tissues ^27–33^.

**Figure 7.**
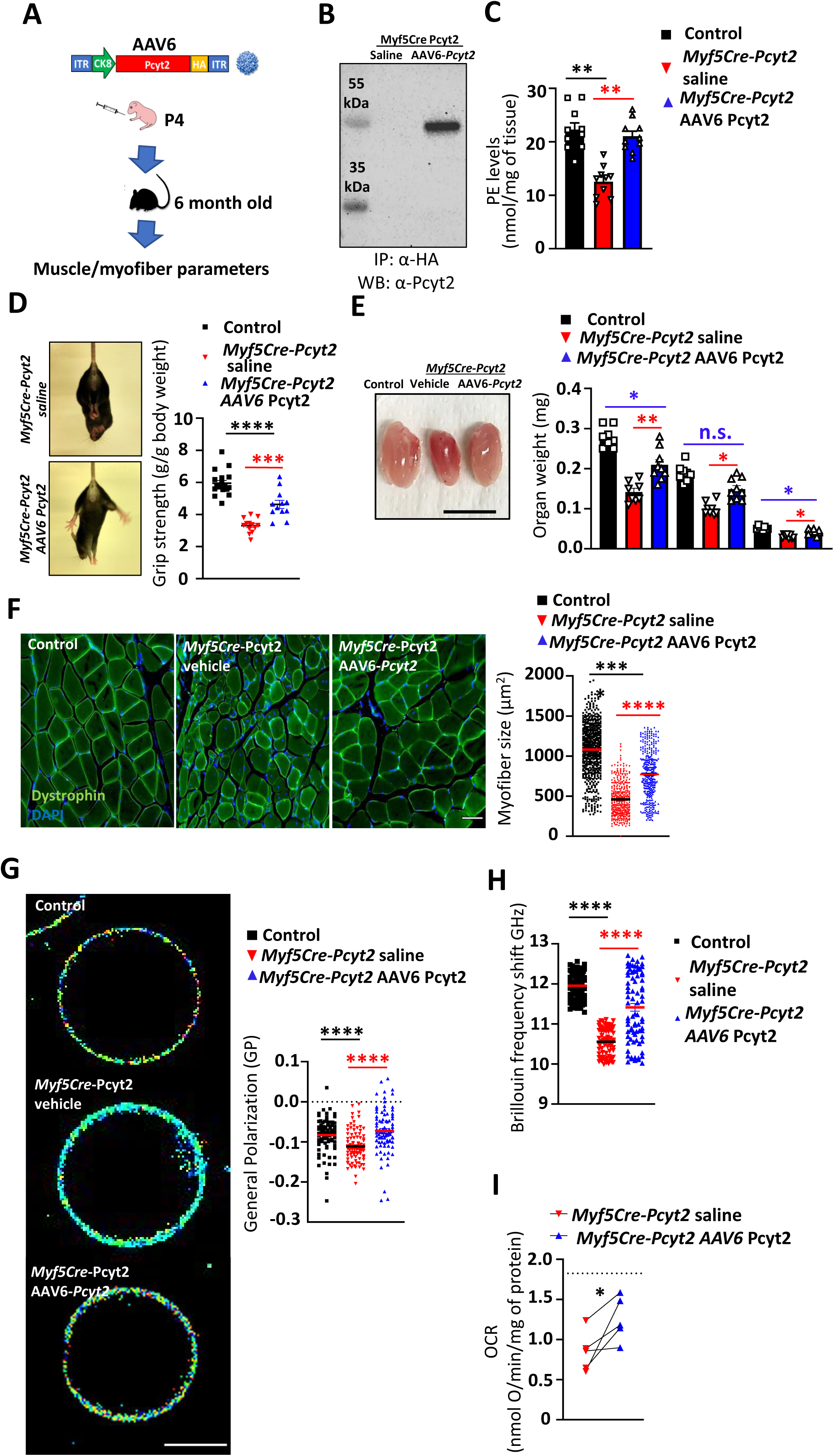
Adenovirus based Pcyt2 gene therapy in mice is efficient for treatment of Pcyt2 deficiency-induced muscle pathology. **(A)** Schematic diagram of muscle specific gene therapy on Pcyt2 deficient background. **(B)** Representative images, and grip strength quantification of control (saline) *Myf5Cre-Pcyt2 (saline) and Myf5Cre-Pcyt2 (AAV6-CK8-Pcyt2HA)* mice. **(C)** Representative appearance and skeletal muscle weight isolated from 6 month old control, *Myf5Cre-Pcyt2 saline* treated *and Myf5Cre-Pcyt2 AAV6-Pcyt2* treated mice. Each dot represents single mice. QA, quadriceps; GC, gastrocnemius; TA, tibialis anterior muscles. Scale bars 1 cm. **(D)** Assessment of Pcyt2HA expression in *Myf5Cre-Pcyt2 AAV6-Pcyt2* treated mice as determined by anti-HA immunoprecipitation, followed by an anti-Pcyt2 blot, from quadriceps muscles isolated 6 months after the gene delivery. **(E)** Total phosphatidylethanolamine levels from quadriceps muscles of control, *Myf5Cre-Pcyt2 saline* treated *and Myf5Cre-Pcyt2 AAV6-Pcyt2* treated mice, isolated 6 months after the treatment. Each dot represents individual mice. **(F)** Representative cross sections and myofiber diameter sizes from 6 month old control, *Myf5Cre-Pcyt2 saline* treated *and Myf5Cre-Pcyt2 AAV6-Pcyt2* mice. Myofibers were imaged using 10X magnification with ≥ 60myofibers analyzed per mouse. n=5 animals per group. Scale bar 100µm. **(G)** Polarization microscopy of NR12S-stained GPMVs isolated from control, *Myf5Cre-Pcyt*2 saline and *Myf5Cre-Pcyt*2 AAV6-*Pcyt2* treated 6 month old mice. GPMVs were isolated from ≥ 100 myofibers from 3 mice per group. Representative images and quantifications are shown. Each dot represents GP values of a single GPMV. Scale bar 5μm. **(H)** Surface stiffness analysis as measured by Brillouin frequency shift (BFS) from isolated myofibers of control, *Myf5Cre-Pcyt*2 saline and *Myf5Cre-Pcyt*2 AAV6-*Pcyt2* treated 6 month old mice. Each data point in the right panel represents a BFS peak value of the myofiber surface. 20 myofibers from n=4 mice per group were analyzed. **(I)** Muscle mitochondrial function of control, *Myf5Cre-Pcyt*2 saline and *Myf5Cre-Pcyt*2 AAV6-*Pcyt2* treated mice as assessed by measurements of complex I linked activity on muscle tissue lysates isolated from 6 months old mice. Each dot represents individual mice, N=5 mice per group. Paired Student t-test was used for statistical analysis. Dashed line indicates the average value of mitochondrial activities measured from 5 individual control mice. Data are shown as means ± SEM. Each dot represents individual mice. *p < 0.05, **p < 0.01, ***p < 0.001, and ****p < 0.0001, n.s. not significant. Unless otherwise indicated, Multiple comparison One-Way ANOVA with Welch correction was used for statistical analysis unless stated otherwise.

**Figure 8.**
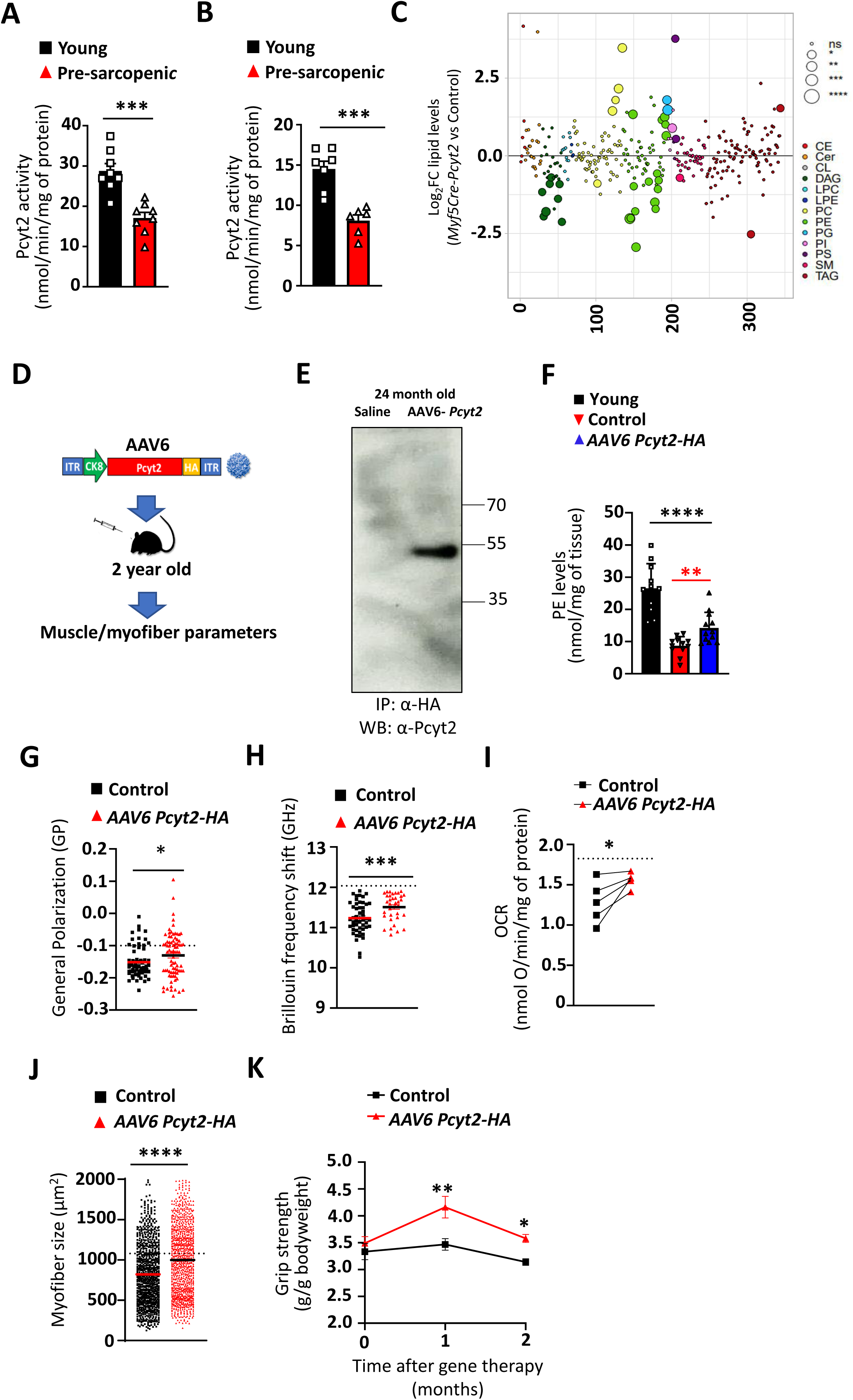
Pcyt2 activity is reduced in aged muscles from humans and mice and Pcyt2 gene delivery ameliorates age-related sarcopenia in sarcopenic mice. **(A)** PCYT2 activity in quadriceps muscle biopsied from young (20-30yr) and middle aged (45-62yr) healthy human volunteers. Each dot represents individual human. Unpaired Student t-test with Welch correction was used for statistical analysis. **(B)** Pcyt2 activity in quadriceps muscles from young (6 month) and pre-sarcopenic (24 month old) C57B6/J mice. Each dot represents individual mouse. Unpaired Student t-test with Welch correction was used for statistical analysis. **(C)** Lipidomics analyses from quadriceps muscles isolated from young (6 month old) and pre-sarcopenic (24 month old) C57B6/J mice. mice. Data are shown relative to control values. CE-cholesterol ester; Cer-Ceramides; DAG-diacylglycerols; LPC-lysophosphatidylcholines; LPE-lysophosphatidylethanolamines; PC-phosphatidylcholines; PE-phosphatidylethanolamines; PG-phosphatidylglycerols; PI-phosphatidylinositols; PS-phosphatidylserines; SM-sphingomyelins; TAG-triacylglycerols. n=5 mice per group. **(D)** Schematic diagram of adenovirus based, muscle specific delivery of Pcyt2 to pre-sarcopenic 24 months old C57B6/J mice. **(E)** Assessment of Pcyt2HA expression as determined by anti-HA immunoprecipitation, followed by an anti-Pcyt2 blot, from quadriceps muscles isolated 2 months after the gene delivery. **(F)** Total phosphatidylethanolamine levels from quadriceps muscles of from young (6 month old), and aged (26 month old) control (saline) and *AAV6-CK8-Pcyt2HA* transduced C57B6/J mice two months after AAV6 injection. Each dot represents individual mice. **(G)** Polarization microscopy of NR12S-stained GPMVs isolated from aged (26 month old) control (saline) and *AAV6-CK8-Pcyt2HA* transduced C57B6/J mice two months after AAV6 injection. GPMVs were isolated from ≥ 100 myofibers from 3 mice per group. Each dot represents GP values of a single GPMV. Scale bar 10μm. Unpaired Student t-test with Welch correction was used for statistical analysis. **(H)** Surface stiffness analysis as measured by Brillouin frequency shift (BFS) from isolated myofibers of aged (26 month old) control (saline) and *AAV6-CK8-Pcyt2HA* transduced C57B6/J mice two months after AAV6 injection. Dashed line indicates the average value of Brillouin frequency shift (BFS) measured from 5 young (6 month) mice. Each data point in the right panel represents a BFS peak value of the myofiber surface. 7 myofibers from n=7 mice per group were analyzed. Unpaired Student t-test with Welch correction was used for statistical analysis. **(I)** Muscle mitochondrial function of aged (26 month old) control (saline) and *AAV6-CK8-Pcyt2HA* transduced C57B6/J mice two months after the treatment as assessed by measurements of complex I linked activity. Each dot represents individual mice. Paired Student t-test was used for statistical analysis. Dashed line indicates the average value of mitochondrial activities measured from 5 individual control mice. **(J)** Myofiber diameter sizes from aged (26 month old) control (saline) and *AAV6-CK8-Pcyt2HA* transduced C57B6/J mice two months after the AAV6 injection. Dashed line indicates the average value of myofiber diameter sizes measured from 5 individual young (6 month old) mice. Myofibers were imaged using 10X magnification with ≥ 180 myofibers analyzed per mouse. n=5 animals per group. Scale bar 100µm. Unpaired Student t-test with Welch correction was used for statistical analysis. **(K)** grip strength quantification of muscle specific gene therapy from aged (26 month old) control (saline) and *AAV6-CK8-Pcyt2HA* transduced C57B6/J mice one and two months after AAV6 injection (n=11-15 per group). Repeated Measures Two-Way ANOVA with Bonferroni correction was used for statistical analysis. Data are shown as means ± SEM. Each dot represents individual mice. *p < 0.05, **p < 0.01, ***p < 0.001, and ****p < 0.0001, n.s. not significant. Unpaired Student t-test with Welch correction was used for statistical analysis unless stated otherwise.

To further explore muscle-specific deletion of Pcyt2, we crossed *Pcyt2^flox/flox^* mice with *Mck-Cre* animals to generate *MckCre-Pcyt2* offspring. Such *MckCre-Pcyt2* mice have been previously reported with an apparent beneficial effect on muscle health^34^, which is in contrast to our findings in *Myf5Cre-Pcyt2* mice and *PCYT2* mutant patients. Critical membrane myotome development is established early in utero, facilitated by addition and fusion of muscle satellite cells (MSC) and adult myoblasts and perturbations in these early events manifest as rapid onset and severe dystrophies ^35,36^. Given that McKCre is active late in muscle formation (peak Cre activity P10), and the mature muscle without affecting the early myotome, MSC and myoblasts^37,38^ ^35,36^, coupled with slow membrane PE turnover^39^, might explain these differences. Importantly, whereas young *MckCre-Pcyt2* mice up to 4 months of age did not display degenerative phenotypes ^34^, older *MckCre-Pcyt2* mice displayed muscle weakness phenotype with a late onset at 18 months of age (Extended Data Figure 8M,N). Overall, these data show that genetic inactivation of *Pcyt2* specifically in muscle results in progressive muscle weakness, muscle wasting, atrophy and a shortened lifespan.

### *Pcyt2* muscle deficiency alters mitochondrial function

Given that phosphatidylethanolamines are abundant in cell membranes but also mitochondrial membranes^40^, we analyzed if *Pcyt2* deficiency affects muscle mitochondrial homeostasis. Parallel to whole skeletal muscle tissue lipidome changes, there was also a significant reduction of PE species in muscle mitochondria from *Myf5Cre-Pcyt2* mice (Figure 4A). Global RNA transcriptome analyses of muscle from 10 day old control and *Myf5Cre-Pcyt2* mice revealed differential enrichment of genes associated with mitochondrial dysfunction in the mutant muscles (Figure 4B). Interestingly, when we analyzed skeletal muscle mitochondrial activity from 2 months old *Myf5Cre-Pcyt2* mice, we observed an increase in mitochondrial activity, followed by a drop in the activity in the muscle mitochondria isolated from 6 months old *Myf5Cre-Pcyt2* mice (Figure 4C,D; Extended Data Figure 9A,B). Further, we observed an increase in mitochondrial ROS in the isolated myofibers, as well as increased levels of anti-oxidant catalase activity and protein oxidative damage in the skeletal muscle of 6 months old Myf5Cre-Pcyt2 mice (Figure 4E-G). As expected, markers of cellular stress (pJNK, Foxo1) and muscle wasting (Atrogin, MuRF1, Fbx031) were increased in the muscles of 6 months old *Myf5Cre-Pcyt2* mice (Figure 4H,I). The ultrastructural morphology and contents of mitochondria appeared unchanged in muscles even in adult mutant mice with apparent phenotype (Extended Data Figure 9C,D). Next, we tested if the accumulation of the Pcyt2 substrate phosphoethanolamine was causing the observed mitochondrial changes. Compared to control mice, there was no additive, inhibitory effect of phosphoethenolamine on the activity of skeletal muscle mitochondria isolated from *Myf5Cre-Pcyt2* mice (Extended Data Figure 9E), suggesting that intra-mitochondrial overaccumulation of phosphoethanolamine is not responsible for the observed reduction in activity of skeletal muscle mitochondria from *Myf5Cre-Pcyt2* mice.

We also tested a potential role of Pcyt2 and Pcyt2-derived PE in sarcolemmal Ca^2+^ handling and autophagy. We failed to observe any structural changes or alterations in Ca^2+^ release and Ca^2+^ uptake in isolated myofibers from 6-month-old *Myf5Cre-Pcyt2* mice manifesting gross phenotypes (Extended Data Figure 10 A-C),suggesting that sarcoplasmic reticulum Ca^2+^ handling was preserved. PE conjugation to ATG8 is also necessary for autophagy ^41^. However, PE-ATG8 conjugation was comparable in both quadriceps and diaphragm muscles of 6-month-old control and *Myf5Cre-Pcyt2* mice under both fed and fasted conditions and there was also no evident accumulation of p62/SQSTM1 under both fed and fasting conditions (Extended Data Figure 10 D-I)Of note, it has been previously shown that genetic inactivation of autophagy and LC3 PE lipidation in Myf5-derived lineages induces brown fat over-activation, but does not lead to muscle weakness, degeneration, not muscle dystrophy ^42^; contrasting with our *Myf5Cre-Pcyt2* mice. These data suggest that loss of Pcyt2 does not affect PE-modifications in autophagy or sarcoplasmic reticulum but alters mitochondrial function.

Given that we observed impaired mitochondrial activity in the skeletal muscle of *Myf5Cre-Pcyt2* mice, we attempted to improve the pathological features of *Myf5Cre-Pcyt2* mice by administering mitochondria-targeted antioxidant tetrapeptide SS-31 daily for two months, starting from 4 months of age. Although there was a mild improvement in the grip strength and skeletal muscle weight, the effect was not significant (Extended Data Figure 9G,H). This suggested that there are additional cellular defects driving the observed pathology due to muscle Pcyt2 deficiency.

### Loss of Pcyt2 directly alters sarcolemmal lipid bilayer organization and rigidity

Impaired sarcolemmal stability causes myofiber degeneration in muscular dystrophies ^43^. Our muscle whole tissue lipidomics data showed a significant decrease in PEs containing long chain fatty acids (FAs)(Figure 1J,K), which are abundant membrane lipids ^40^. Therefore, we hypothesized that the reduced abundance of PEs containing long chain FAs in *Myf5Cre-Pcyt2* mice might also affect formation and stability of the sarcolemmal lipid bilayer, driving the muscular pathology. To test this hypothesis, we first evaluated whether the organization of the sarcolemmal lipid bilayer was altered in *Myf5Cre-Pcyt2* mice. Spectral imaging of NR12S-stained giant plasma membrane vesicles (GPMVs) provides structural information of lipid bilayers in their native compositional complexity, which enables measurements of membrane packing, affecting bending rigidity ^44^. We derived the parameter of general polarization (GP), with higher values corresponding to tightly packed, rigid lipid bilayer and lower values corresponding to loosely packed, and soft bilayer. Strikingly, polarization microscopy of GPMVs derived from *Myf5Cre-Pcyt2* myoblasts displayed loosely packed and less rigid lipid bilayer as compared to control myoblasts derived GMPVs (Figure 5A-B). To further address if the membrane bilayer changes persist in terminally differentiated myofibers, we assessed GPMVs from myofibers immediately after isolation from the tissue (Figure 5C). GPMVs isolated from myofibers of *Myf5Cre-Pcyt2* mice again showed significant reduction of lipid packing and soft lipid bilayer (Figure 5D). We infer that lipid bilayer membrane organization and rigidity are disrupted in both myoblasts and terminally differentiated myofibers that lack Pcyt2.

To directly address how these structural-chemical changes of the membrane lipid bilayer affect mechanical properties of the whole myofibers, we employed high-resolution Brillouin Light Scattering Microscopy (BLSM) on isolated myofibers; BLSM is used to assess the elastic modulus of bio-materials ^45^. Scans using BLSM revealed a significant reduction in the surface stiffness of myofibers isolated from *Myf5Cre-Pcyt2* mice compared to controls (Figure 5E-F). Atomic force microscopy on single myofibers further confirmed that *Myf5Cre-Pcyt2* myofibers have a higher degree of membrane deformity after applying pressure at a nanoscale level and reduced membrane stiffness compared to *Pcyt2*-expressing muscle cells (Figure 5G,H). Thus, loss of *Pcyt2* in skeletal muscle fibers results in an altered architecture of membrane lipid bilayers, directly perturbing sarcolemmal lipid bilayer mechanical properties of rigidity and stiffness.

### *Pcyt2* deficiency leads to impaired sarcolemmal integrity and strain intolerance

An intact cell membrane architecture is critical for membrane barrier function. In particular, the sarcolemma undergoes recurrent injury, for example via mechanical strain such as during exercise, and needs to be repaired for proficient skeletal muscle function ^46^. To determine if the perturbed architecture of the sarcolemma in *Myf5Cre-Pcyt2* mice leads to altered permeability, we injected 6-month-old control and *Myf5Cre-Pcyt2* mice intraperitoneally with Evans blue (<1 kDA) ^47^. We observed an extensive accumulation of Evans blue in the quadricep muscles of *Myf5Cre-Pcyt2* mice relative to controls (Figure 6A,B To further explore sarcolemmal stability, we induced laser mediated membrane microinjury on freshly isolated myofibers and quantified the extent of damage in real time, by measuring intracellular entry of the dye fm1-43, a styryl dye that is nonfluorescent in aqueous solution (extracellular) but increases its fluorescence after intracellular entry and binding to lipid vesicles ^48^. Control myofibers displayed minimal influx of fm1-43 following laser injury, whereas *Myf5Cre-Pcyt2* myofibers showed increased permeability to fm1-43 in response to the laser microinjury (Figure 6C; Video S1-S4). Thus, loss of Pcyt2 results in compromised myofiber membrane permeability *in vitro* and *in vivo*.

To directly address sarcolemmal durability to strain *in vivo*, we subjected littermate control and *Myf5Cre-Pcyt2* mice to a downhill running regimen on a treadmill; such eccentric exercise is a potent inducer of sarcolemma strain, while having very low energy requirement compared to concentric exercise of the same load ^49^. During the early acclimatization phase (low speed downhill running; 4 meters min^-1^ for 40 min) and the intermediate phase (4 meters min^-1^ for 40 min plus 9 meters min^-1^ for 20 minutes) of the eccentric exercise, *Myf5Cre-Pcyt2* mice performed similarly to their littermates (Figure 6D). However, during the late stress phase (20 meters min^-1^ for 20 minutes), *Myf5Cre-Pcyt2* mice failed to complete the exercise (Figure 6D; Video S5). We therefore analyzed the skeletal muscle and after the last phase of training. The weights of the quadricep and gastrocnemius muscles increased after training in control mice. In contrast, the muscles of *Myf5Cre-Pcyt2* mice failed to undergo physiologic hypertrophy (Extended Data Figure 11A); instead their quadriceps muscle exhibited foci of inflammatory cell infiltrates, ectopic fatty cell deposits and fibrosis, resulting in a marked overall myositis score (Figure 6E-F, Extended Data Figure 11B). In parallel, we observed a significant increase in blood muscle creatine levels after exercise (Figure 6G). Immunostaining for Dysferlin, a protein critical to sarcolemmal membrane repair, revealed that Dysferlin was aberrantly localized in *Myf5Cre-Pcyt2* quadricep muscles after training (Extended Data Figure 11C). In addition, we observed disorganized F-actin networks in *Myf5Cre-Pcyt2* mice after training (Figure 6H). Of note, the number of Pax7^+^ progenitors in the quadricep muscle appeared similar between age-matched trained control and *Myf5Cre-Pcyt2* mice (Extended Data Figure 11E). Thus, Pcyt2 PE synthesis is required for sarcolemmal integrity under strain, preventing excessive muscle damage and enabling physiologic hypertrophy during eccentric exercise.

### Muscle-specific *Pcyt2* gene therapy rescues muscle weakness in *Pcyt2* mutant mice

Currently, there is no treatment for the disease caused by *PCYT*2 deficiency. Since gene therapies have made great advancements and shown promise in the treatment of rare diseases ^50,51^ we sought to therapeutically ameliorate the muscle weakness *Myf5Cre-Pcyt2* mice using muscle specific delivery of HA-tagged Pcyt2. Briefly, we cloned mouse Pcyt2 under the control of the muscle-restricted creatine kinase 8 (CK8) promoter/enhancer into an AAV-based gene delivery vector, in this case AAV6 ^52,53^. Of note, AAV6 based gene therapies are currently in clinical trials for Duchenne muscular dystrophy ^53^. The *AAV6:CK8:Pcyt2-HA* vector or saline (as control) were injected into 4 day old *Myf5Cre-Pcyt2* mice and the muscles of these mice were assessed 6 months after the treatment (Figure 7A). This approach resulted in Pcyt2-HA protein overexpression as well as increased PE levels in the skeletal muscles of *Myf5Cre-Pcyt2 AAV6:CK8:Pcyt2-HA* injected mice compared to untreated *Myf5Cre-Pcyt2* controls (Figure 7B,C, Extended Data Figure 12A).

Strikingly, the *AAV6:CK8:Pcyt2-HA* treated mutant mice displayed a significant increase in grip strength, increased skeletal muscle mass and myofiber diameter (Figure 7D-F; Extended Data Figure 12B). Moreover, skeletal muscles from *Myf5Cre-Pcyt2 AAV6:CK8:Pcyt2-HA* injected mutant mice, as compared to their untreated littermates, displayed improved muscle membrane parameters (Figure 7G,H) and improved mitochondrial respiration (Figure 7I). Thus, muscle specific AAV6-based gene delivery of wild type *Pcyt2* can efficiently ameliorate the muscle pathology of *Myf5Cre-Pcyt2* mice.

### Muscle-specific *Pcyt2* gene therapy improves muscle strength in aging

Sarcopenia and progressive muscle atrophy are critical determinants of frailty in aging. Muscle aging is commonly associated with diminished membrane integrity, increased susceptibility to damage, and diminished repair after exercise ^54–61^. As *Myf5Cre-Pcyt2* mice displayed degenerative features that are also found in aging muscles, we assessed a potential role of *Pcyt2* in muscle aging. Indeed, Pcyt2 mRNA expression and enzymatic activity in quadricep muscle were reduced in aged, pre-sarcopenic mice compared to young mice (Figure 8A, Extended Data 12B). Importantly, PCYT2 activity and levels were substantially decreased in quadricep muscle biopsies of otherwise healthy 45-62 year-old compared to 20-30 year-old humans (Figure 8B, Extended Data 12C,D). Thus, in mice and humans, Pcyt2 expression and even more activity decline with aging. Accoupling this, we found that in the muscles of aged, pre-sarcopenic mice, some of the most significantly affected lipids were PE species (Figure 8C). To test whether increasing the levels of Pcyt2 can improve muscle function in aged mice, we aimed to rejuvenate aged muscles via overexpression of *Pcyt2*. *AAV6:CK8:Pcyt2-HA* or saline (as control) were injected retro-orbitally into 24 month-old male C57B6/J mice (Figure 8D). and the expression of Pcyt2-HA and increase of total muscle PE levels in the quadricep muscle was confirmed after 2 months (Figure 8E,F). Using our newly established microscopy-based analysis pipeline, we addressed whether the observed *AAV6:CK8:Pcyt2-HA* dependent increase of PE lipids in the muscles of aged mice would also reflect on myofiber membrane physical parameters. Indeed, there was a significant increase in muscle membrane stiffness as measured by both polarization microscopy and Brillouin spectroscopy (Figure 8G,H). Moreover, the bioenergetics of the muscle showed beneficial changes, with improved mitochondrial capacity and improved respiratory control ratio, which is a measure of ATP production efficiency (Figure 8I, Extended Data Figure 12E). Accompanying this, there was a significant increase in the myofiber diameter of *AAV6:CK8:Pcyt2-HA* mice compared to control mice 2 months after gene delivery (Figure 8J).

Remarkably, we observed significantly improved grip strength at 1 and 2 months after gene delivery in *AAV6:CK8:Pcyt2-HA* mice compared to control mice (Figure 8E, Extended Data Figure 12F). Thus, Pcyt2 overexpression has a beneficial effect on muscles in aged mice.

In summary, our results uncover a critical and conserved role for Pcyt2 and Pcyt2-regulated lipid biosynthesis. We show that loss of Pcyt2-dependent lipid biosynthesis causes a previously unrealized form of muscular dystrophy, characterized by an aberrant muscle development, progressive muscle weakness and wasting, failure to thrive and shortened lifespan. Our work reveals that with *Pcyt2* deficiency, reduction of long chain PE synthesis compromises the sarcolemma lipid bilayer stability, as well as myofiber mitochondria homeostasis. Given that PEs are predominantly membrane lipids, we infer that Pcyt2-dependent PE synthesis is essential for the lipid bilayer of sarcolemma and PE enriched membranes of mitochondria.

This form of muscular dystrophy is very rare, in that a lipid species provides mechanical support to cellular membranes, as opposed to other forms of dystrophies that are caused by aberrations of cytoskeletal, mitochondrial or extracellular proteins^24^, and may thus also have distinct therapeutic implications.

Whereas muscular dystrophy is typically caused by the disruption of proteins that support the sarcolemma, we show that loss of *Pcyt2* leads to intrinsic changes of the membrane lipid bilayer, thus representing a unique disease mechanism.

Besides being essential to sarcolemmal lipid bilayer composition and stability, PE lipids are also abundant in the inner mitochondrial membrane. Although majority of mitochondrial PEs are derived via decarboxylation of phosphatidylserine inside the mitochondria ^62^, the observed mitochondrial dysfunction with increased generation of mtROS in myofibers of Myf5Cre-Pcyt2 mice clearly demonstrate that Pcyt2-dependent PE synthesis is additionally important in generating mitochondrial PE lipids. Decreased mitochondrial membrane viscosity coupled with impaired mitochondrial activity was already observed in degenerative diseases ^63^, while decreased membrane viscosity increases ROS diffusion across the bilayer ^64,65^. This phenomenon could explain alterations of mitochondrial respiration as well as increase in ROS levels in myofibers which coupled with sarcolemmal instability, triggers muscle mitochondrial functional decline, myofiber stress, increased ROS levels, oxidative protein damage and activation of the JNK-FoxO1 axis of muscle atrophy in Myf5Cre-Pcyt2 mice. Consequently, these perturbations result in dramatic muscle degeneration, progressing to a severe dystrophy with inflammation and shortening of lifespan.

Intriguingly, we and others^34^ have observed an initial increase of mitochondrial activity in Pcyt2 deficient muscles, without an increase in total mitochondrial numbers. This initial response most likely represents a compensatory response typical of mitochondrial diseases^66^.

Mouse models that initiate gene loss very early in myotome development, faithfully recapitulate human muscular dystrophies in many pathological features including stunted growth and progressive muscle degeneration with shortened lifespan ^24,35,36^. MckCre-Pcyt2 deficient mice did not display muscular weakness at an early adulthood ^34^, and we observed muscle weakness in old (20 months old) MckCre-Pcyt2 mice. Mck promoter drives Cre expression late in the muscle development (peak activity P10)^37^. However, critical muscle membrane organization occurs very early in the developing myotome^24^. Thus relatively later Cre activation via Mck promoter would miss this critical developmental stage, which is not the case for inherited, disease-causing mutations in humans. Moreover, given that MckCre activity is restricted to mature muscles ^37,38^, this bypasses the muscle stem cell pool rendering them “wild type”, which can then actively repopulate and repair the Pcyt2 deficient muscle, and delay muscle degeneration. This phenomena of dependence of disease severity on the timing of the gene disruption was already observed for several animal models of membrane-related human dystrophies, where only mouse mutants with early gene deletion faithfully recapitulated patient phenotypes ^35,36^. Our findings in zebrafish model, mouse mutant and aged models, as well as in rare disease patients with weakness and muscle wasting ^67^ support that the Pcyt2-dependent Kennedy pathway is essential for muscle health.

As seen in rare disease patients, failure to thrive preceded the apparent progressive weakness and degeneration in Myf5Cre-Pcyt2 mice. Muscle growth by myoblast fusion and hypertrophy is essential for both muscle and whole-body size ^68^. Both mechanisms of muscle growth were affected in Myf5Cre-Pcyt2 mice. Apart from being important membrane building blocks, phosphatidylethanolamines are important in modulating membrane physicochemical properties ^4^. Due to their relatively small polar head group and consequently a conical shape ^4^, phosphatidylethanolamines form a negative membrane curvature required for membrane bending during the cell fusion process^69^. Therefore, any genetic insufficiency of phosphatidylethanolamines would negatively affect the efficiency of both membrane neo-genesis during tissue growth and cellular fusion, affecting tissue growth. However, impaired growth as well as myoblast fusion wouldn’t necessarily result in degenerative changes later on in adulthood ^70–73^, while several long lived mouse models are also smaller ^74^.

Our findings indicate that the muscle tissue is especially vulnerable to loss of Pcyt2 and Pcyt2 synthesized PE. It is well established that distinct tissues have a diverse membrane lipid composition ^4^ and may be differentially dependent on Pcyt2. Indeed, mining the Achilles Depmap data portal, which contains gene essentiality scores from 769 cell lines ^75^, we found that Pcyt2 is not essential for a large majority of the tested cell lines (4.8% dependent cell lines). For comparison, many more cell lines (54% dependent cell lines) are dependent on choline-phosphate cytidylyltransferase (Pcyt1a), a bottleneck enzyme for synthesis of phosphatidylcholines in the parallel branch of the Kennedy pathway. The muscle dependency on Pcyt2 derived PE might be explained by the general chemical properties of PE lipids. Increasing PE concentrations increase the viscosity of the liposomes ^76^, therefore we hypothesize that the constant mechanical strain and contraction of the myofibers render muscle membranes dependent on PE for mechanical support. The essential dependency of myofibers on Pcyt2 derived PE compared to other cell types, is supported by our findings from various tissue-specific mutants. Moreover, current pathophysiological symptoms in the rare disease patients are mainly restricted to growth and neuromuscular parameters. This indicates that other cell types are able to engage alternative molecular mechanisms to compensate for the deficiency in Pcyt2-dependent PE synthesis. Future research should illuminate which synthesis or lipid uptake mechanisms are responsible for this. Interestingly, inherited mutation in choline kinase beta (Chkb) results in loss of synthesis of another membrane lipid, phosphocholine and consequently phosphatydilcholine, causing neuronal pathology as well as muscular dystrophy without affecting other tissues^77,78^, thus resembling certain pathological features of inherited PCYT2 mutations. It is intriguing that nervous and muscle tissue are particularly vulnerable to deficiency of certain membrane lipids as opposed to other cell types stimulating a wider field of research on cell type-specific changes in cell membrane lipid composition and lipid bilayer physicochemical parameters in various biological processes and pathological conditions.

Muscle loss and sarcopenia are critical hallmarks of aging, and a leading cause of frailty and dependency. We found that Pcyt2 levels and activity markedly declined in muscles from aged rodents and humans. Of note, we chose our time-point for this analysis at a stage that is pre-sarcopenic to avoid secondary effects of frailty. Decreased expression of Pcyt2 mRNA was recently observed in aged rat muscles ^79^. It is possible that this reduction occurs as a consequence of a metabolic switch in aged muscle, which in aging appears to be more directed towards triglyceride and cholesterol synthesis ^80^. Indeed, low density lipoprotein, cholesterol oxysterols or LXR (liver X receptor, a transcriptional regulator of cholesterol, fatty acid, and glucose homeostasis) regulate and inhibit Pcyt2 ^81^. Importantly, our data show that Pcyt2 and long chain PE membrane lipids are critically involved in muscle health. Muscle specific AAV based Pcyt2 gene therapy ameliorated the muscle weakness in mutant mice, thus paving the way for treatment of the severe human disease developing as a result in *PCYT2* mutation. Strikingly, we also found that increasing Pcyt2 expression in aged mice improves several parameters of aged myofibers through increasing tissue PE lipid levels, consequently ameliorating muscle strength decline. Given the decline of Pcyt2 in aging, decrease of tissue PE levels in aged muscles, and the beneficial effect upon Pcyt2 gene delivery into aged mice, Pcyt2 upregulation could be considered as a potential treatment to improve muscle frailty, a critical issue in our aging world.

## Acknowledgements

We would like to thank all members of our laboratories for helpful discussions and Life Science Editors for editorial support. We are grateful to Vienna Biocenter Core Facilities: Mouse Phenotyping unit, Histopathology unit, Bioinformatics unit, Biooptics unit, Electron microscopy unit and Comparative medicine unit. We are grateful to the Lipidomics facility and K. Klavins T. Hannich at the CeMM Research Center for Molecular Medicine of the Austrian Academy of Sciences for assistance with lipidomics analysis. We would also like to thank to Tao Huan and Alyssa Hui (UBC Vancouver, Canada) for mouse tissue and mitochondria lipidomics analysis. We thank A. Klymchenko (Laboratoire de Bioimagerie et Pathologies Université de Strasbourg, Strasbourg, France) for providing the NR12S probe. We are thankful to the Sen. Paul D. Wellstone Muscular Dystrophy Cooperative Specialized Research Center Viral Vector Core facility (Seattle, US) for AAV6 production. We would also like to thank Kevin P Campbell and Mary E Anderson (University of Iowa, Carver College of Medicine, Iowa US) for advice on muscle tissue handling. J.M.P. is supported by IMBA, an ERC Advanced Grant, a Wittgenstein award, the T. von Zastrow foundation, and a Canada 150 Research Chair in functional genetics. R.Y. is supported by a PhD studentship from the Wellcome Trust (203995/Z/16/Z). J.S.C. is supported by grants RO1AR44533 & P50AR065139 from the US National Institutes of Health. C.K. is supported by a grant from the Agence Nationale de la Recherche (ANR) (ANR-18-CE14-0007-01). .AV.K. is supported by European Union’sHorizon 2020 research and innovation programme under theMarie Skłodowska-Curie grant agreement No 67544, an Austrian Science Fund (FWF) No P-33799, AW is supported by Austrian Research Promotion Agency (FFG) project No 867674. E.S. is supported by a SciLifeLab fellowship and Karolinska Institutet Foundation Grants. Work in the GSF laboratory is supported by the Austrian Academy of Sciences, the European Research Council (ERC AdG 695214 GameofGates) and the Innovative Medicines Initiative 2 Joint Undertaking (grant agreement No 777372, ReSOLUTE).

## Author Contributions

D.C. together with J.M.P. designed and supervised the mouse study, and wrote the manuscript with the input from the co-authors. All experiments were performed and established by D.C. with the following exceptions: K.E. performed Brillouin light scattering microscopy measurements. E.S. performed GPMVs isolation and image analysis with assistance from D.C. E.K. performed synergic ablation assay and analysis under supervision form Z.R. T.F. performed treadmill experiment with assistance from D.C. and under supervision from Z.R. R.Y. collected and analyzed zebrafish models under supervision from M.L. L.X.H. and V.S. performed lipidomics analysis under supervision of G.S.F. A.T. and S.G. performed Pcyt2 enzyme activity analysis under supervision of M.B. A.A. performed *in vivo* brown fat activity and Ucp1 RT-PCR analysis under supervision from C.K. A.W. performed respiration analysis under supervision from A.K. C.K. and C.S. performed myofiber calcium kinetics experiment, with analysis and supervision under J.V. S.J.F. assisted with western blot experiments. M.N. performed bioinformatic analysis of efficiency of Pcyt2 deletion in mice. A.K. performed myositis scoring. N.D.M. performed atomic force microscopy measurements and data analysis with assistance from D.C. T.S. and B.H. assisted in histological analysis. L.H. performed AAV6 i.v. injections. A.H. assisted in tissue sampling for western blot experiments. S.J. provided *Pcyt2* floxed mice. E.R. and T.G. collected human muscle biopsies. J.S.C. generated AAV6 vector and provided guidance with all AAV experiments. J.M. and S.B. identified PCYT2 human mutant carriers, collected growth data and generated growth curves.

## Extended Data Figure legend

**Extended Data Figure 1.**
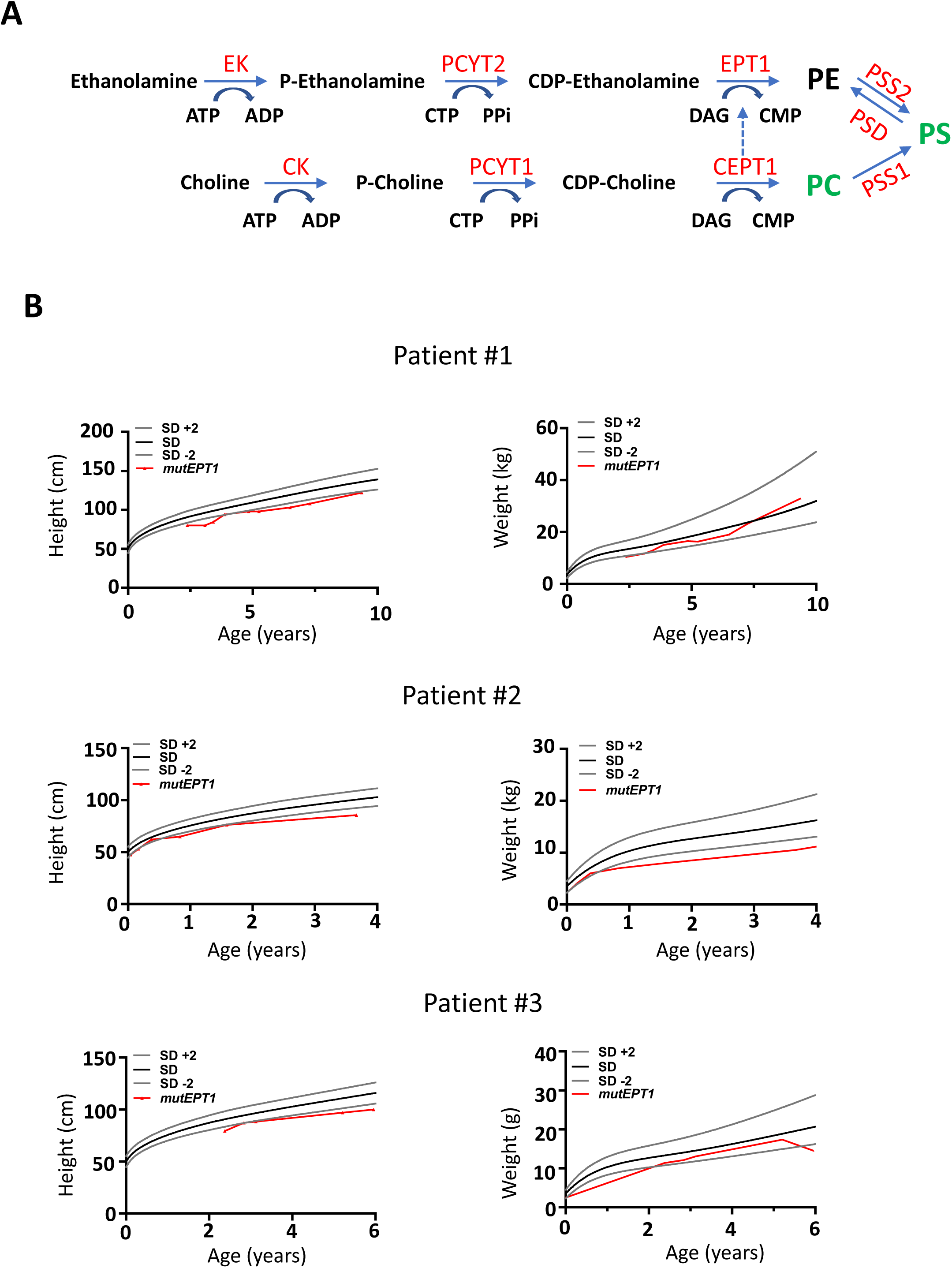
**PE synthesis pathways and *EPT1* rare disease mutation carriers.** **(A)** Schematic diagram of phosphatidylcholines (PC), phosphatidylethanolamines (PE) and phosphatidylserine (PS) phospholipids synthesis. EK-Ethanolamine kinase; PCYT2-CTP:phosphoethanolamine cytidylyltransferase; EPT1-ethanolaminephosphotransferase 1; PSS2-Phosphatidylserine Synthase 2; PSD-Phosphatidylserine decarboxylase; CK-Choline kinase; PCYT1-Choline-phosphate cytidylyltransferase; CEPT1-Choline/ethanolaminephosphotransferase 1; PSS1-Phosphatidylserine Synthase 1. **(B)** Height and weight gains of three patients carrying the homozygous missense variant c.335 G>C (p.Arg112Pro) in the *EPT1* gene. Controls indicate WHO standards of median weights and heights at the respective ages +/-2 standard deviations (SD).

**Extended Data Figure 2.**
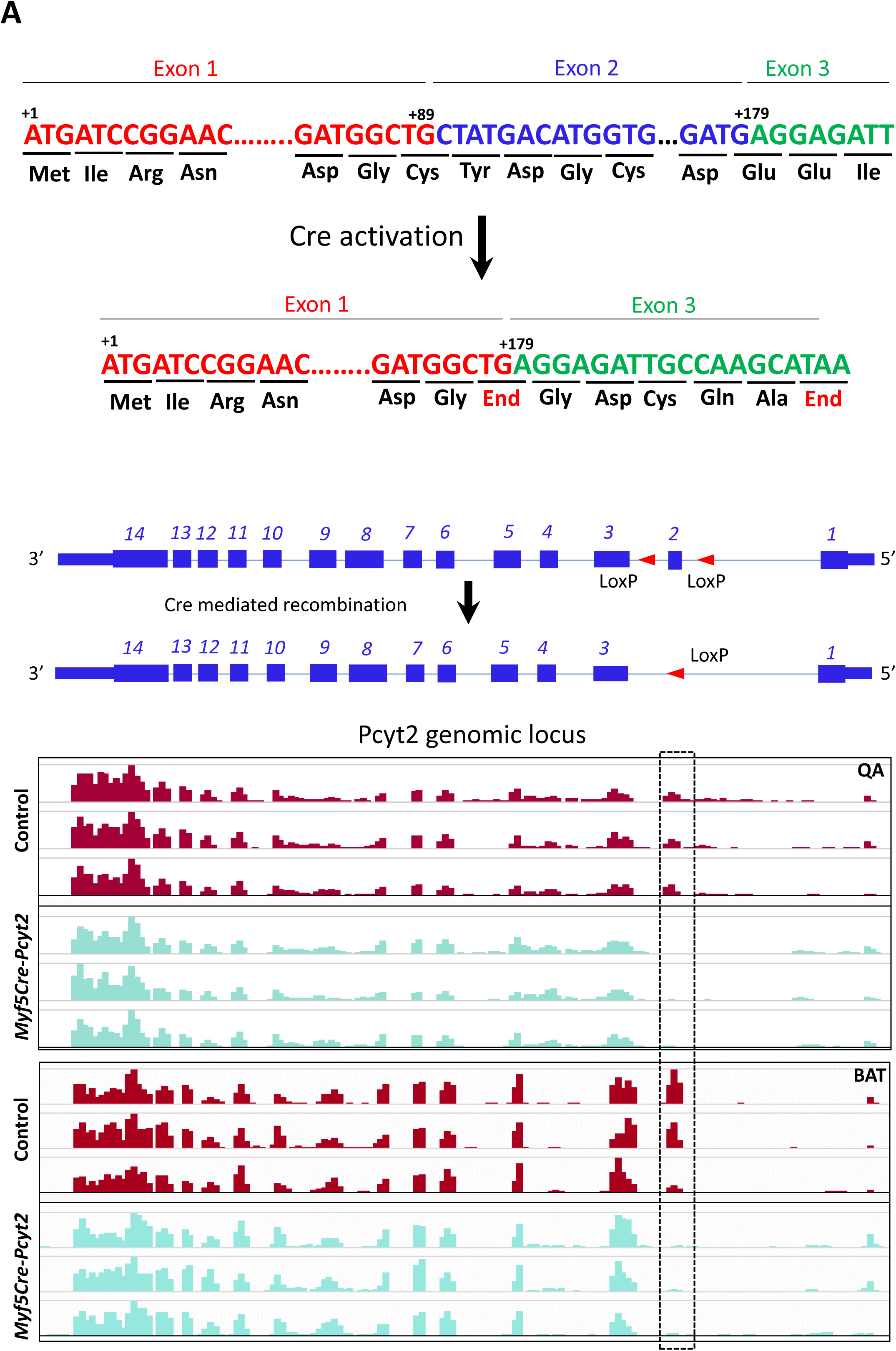
**Analysis of *Pcyt2* deletion in mice.** **(A)** Schematic diagram of exon 2 deletion in *Myf5Cre-Pcyt2* mice and confirmation by RNA sequencing. Exon and introns structures as well as LoxP sites targeted to exon 2 and loss of exon 2 upon Cre-mediated recombination are shown for the murine Pcyt2 locus. n=3 animals per group.

**Extended Data Figure 3.**
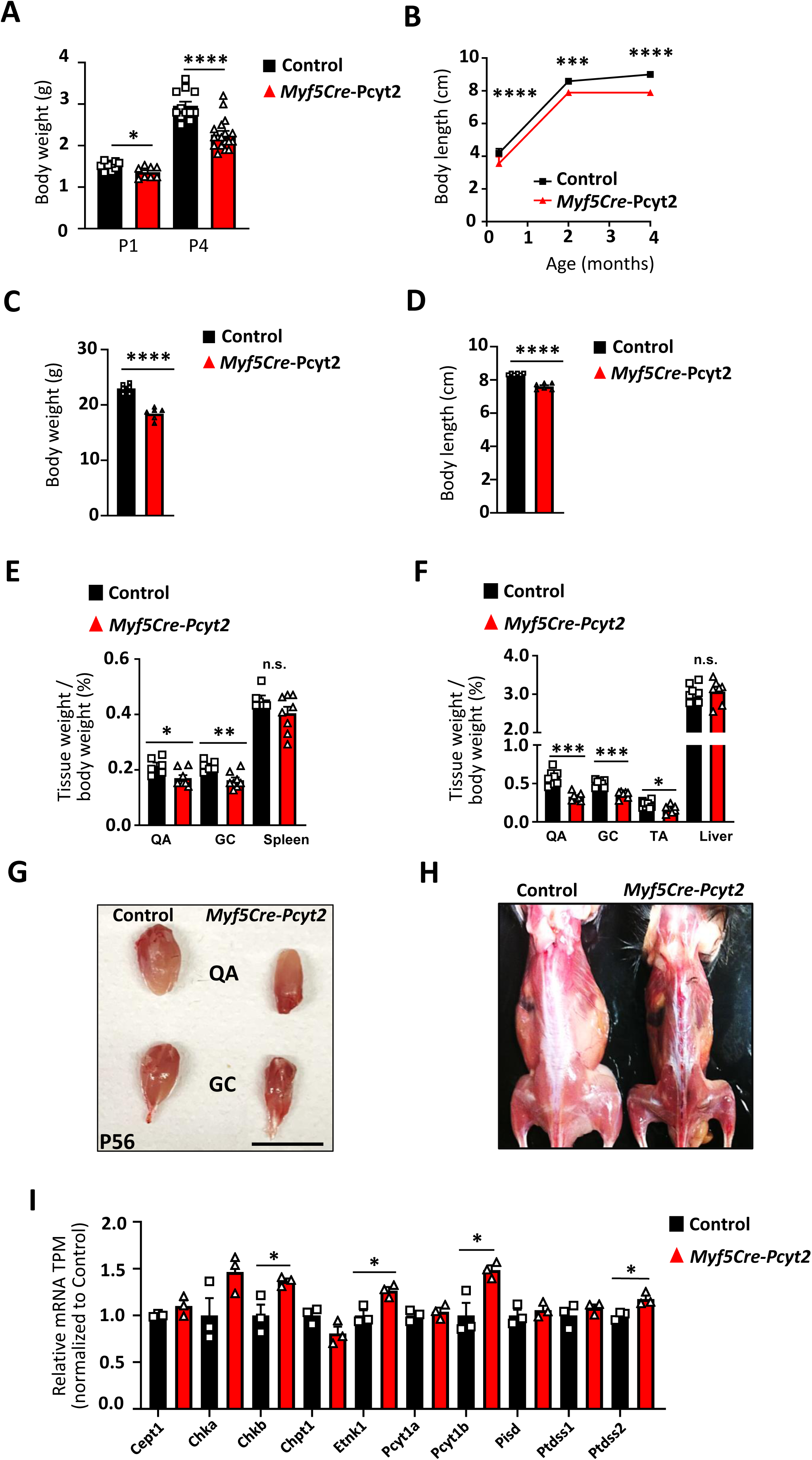
**Characterization of *Myf5Cre-Pcyt2* mice.** **(A)** Body weights of control and *Myf5Cre-Pcyt2* mice at P1 and P4. **(B)** Body length gains of control and *Myf5Cre-Pcyt2* mice. n=6 per group for body length analysis. **(C)** Body weights of 2 months old control and *Myf5Cre-Pcyt2* female mice. **(D)** Body lengths of 2 months old control and *Myf5Cre-Pcyt*2 female mice. **(E)** Skeletal muscle and tissue weight isolated from (E) 10 day and (F) 2 month old (P56) control and *Myf5Cre-Pcyt2* littermate mice. QA, quadriceps; GC, gastrocnemius; TA, tibialis anterior muscles. Liver and spleen weights are shown as controls. n = 6-8 per group Scale bars 1 cm. **(G-H)** Gross skeletal muscle appearance of 56 days old control and *Myf5Cre-Pcyt2* littermates. Data are shown as means ± SEM. Each dot represents individual mice. *p < 0.05, **p < 0.01, ***p < 0.001, and ****p < 0.0001, n.s. not significant (unpaired Student t-test).

**Extended Data Figure 4.**
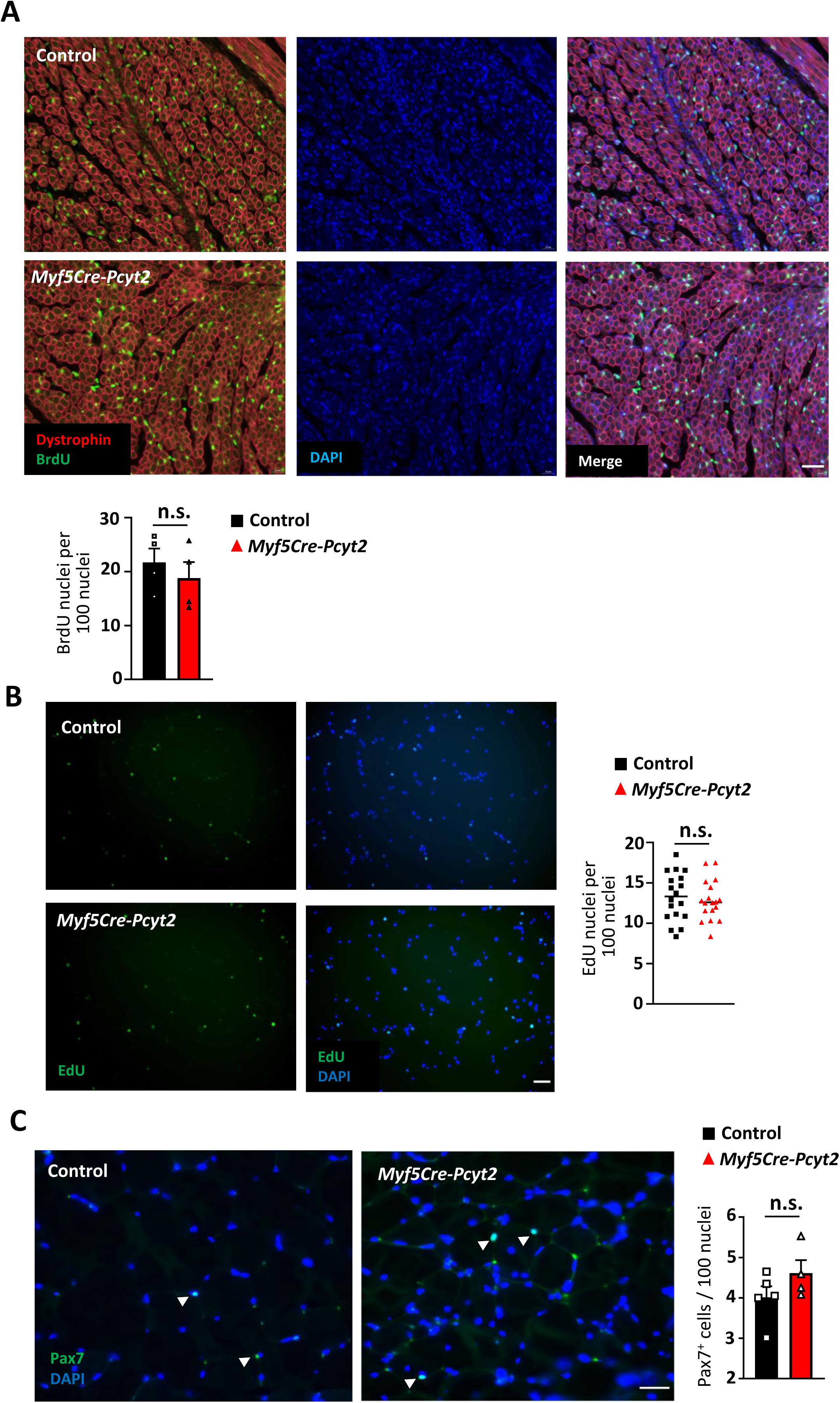
**Myoblast proliferation assessment in *Myf5Cre-Pcyt2* mice.** **(A)** Representative images and quantification of BrdU labeled quadriceps from 2 days old control and *Myf5Cre-Pcyt2* mice. Images were taken under 5x magnification, and ≥2000nuclei were counted and analyzed. N=4 animals per group. Scale bar 60µm. **(B)** Representative images and quantification of EdU labeled primary myoblasts in cell culture isolated from control and *Myf5Cre-Pcyt2* mice. Images were taken under 5x magnification, 6 replicates were analyzed per each cell culture, with ≥100 nuclei counted per replicate. N=3 mice per group. Scale bar 50µm. **(C)** Number of Pax7 positive nuclei in quadriceps from 6 months old control and *Myf5Cre-Pcyt*2 mice. Sections from 3 mice were examined, with ≥100 nuclei per section counted. Scale bar 50µm.

**Extended Data Figure 5.**
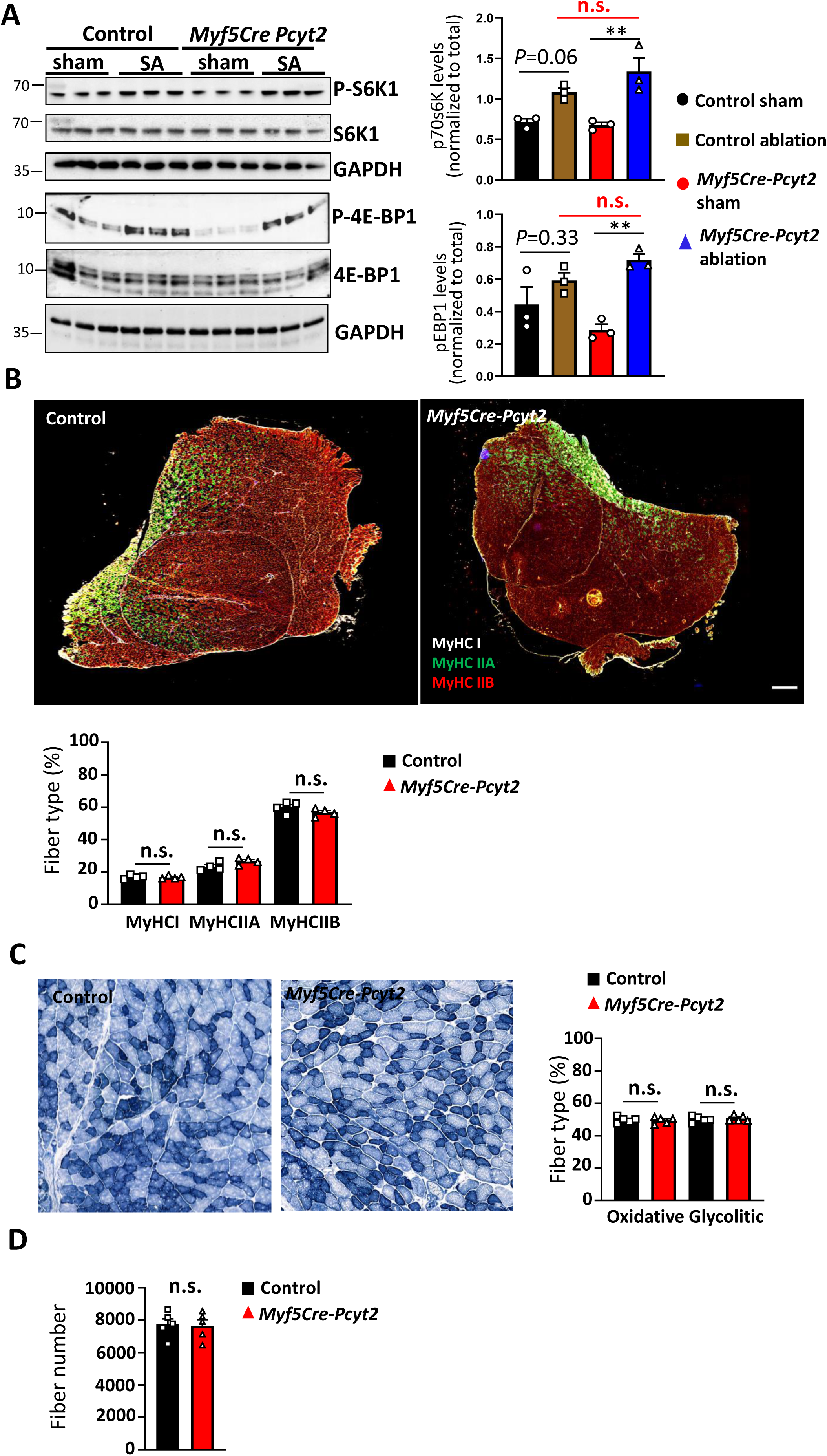
**Myofiber type distribution in skeletal muscle of *Myf5Cre-Pcyt2* mice.** **(A)** Western blot analysis of critical regulators of protein translation RPS6 and eIF4A in overloaded M. plantaris from Control and Myf5Cre-Pcyt2 mice. Each lane represents individual mice. **(B)** Representative images and quantification of MyHC!, MyhCIIA and MyHCIIB fibers in skeletal muscle (quadriceps) from 6 months old control and *Myf5Cre-Pcyt2* mice. Images were taken under 5x magnification, and ≥100 myofibers were counted at 3 different matching histological areas. N=4 animals per group. Scale bar 500µm. **(C)** Representative images and quantification of oxidative and glycolytic fibers in skeletal muscle (quadriceps) from 6 months old control and *Myf5Cre-Pcyt2* mice. Images were taken under 10x magnification, and ≥1000 myofibers were counted at matching histological areas. N=5 animals per group. Scale bar 500µm. (D) Total number of fibers in skeletal muscle (quadriceps) from 6 months old control and *Myf5Cre-Pcyt2* mice. Images were taken under 2.5x magnification. N=4 animals per group. Scale bar 500µm.

**Extended Data Figure 6.**
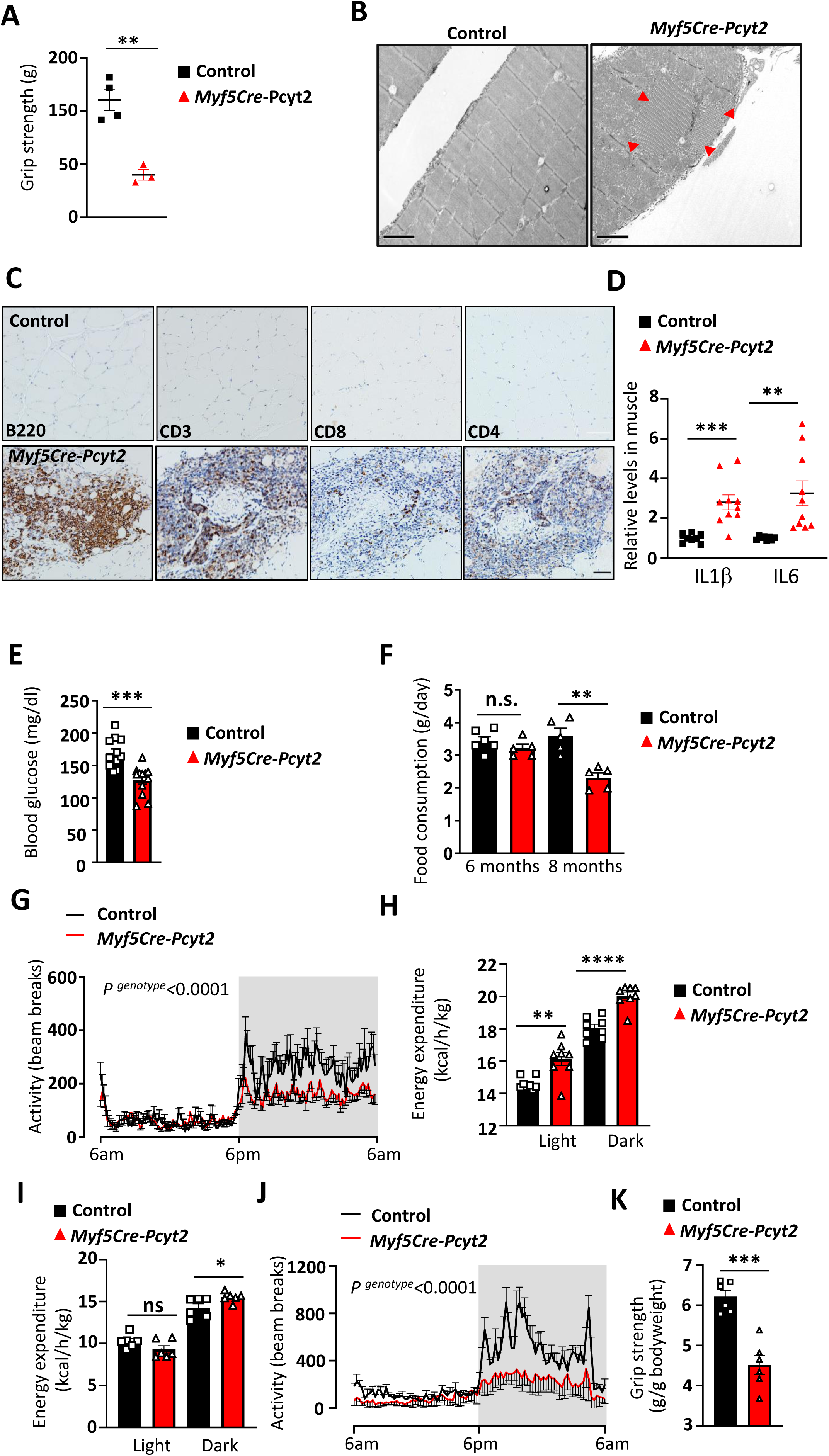
**Muscle inflammation and metabolic assessment of *Myf5Cre-Pcyt2* mice.** **(A)** Representative electron microscopy images of quadriceps of 15 months old control and *Myf5Cre-Pcyt2* mice. Note accumulation of tubular aggregates in the mutant animals (red arrows). Scale bar 2µm. **(B)** Characterization of muscle inflammation in 12 months old *Myf5Cre-Pcyt2* mice. Helper T cells (CD4+) and cytotoxic T cells (CD8+) are shown. Scale bar 100µm for H&E stained and 50 µm for immune cell staining. **(C)** Inflammatory cytokine levels in the quadriceps of 12 months old *Myf5Cre-Pcyt2* mice. **(D)** Fed blood glucose levels on normal chow diet of 8 months old control and *Myf5Cre-Pcyt2* mice. **(E)** Food consumption analysis of 6 month and 8 month old control and *Myf5Cre-Pcyt2* mice **(F)** Cage activity of 6 months old control and *Myf5Cre-Pcyt2* mice (n=12 per group). Multiple ANOVA was used to analyze the data. **(G)** Energy expenditure of 6 months old control and *Myf5Cre-Pcyt2* mice during the resting (light) and active (dark) phases. **(H)** Cage activity under thermoneutrality of 6 months old control and *Myf5Cre-Pcyt2* mice (n=6 per group). Multiple ANOVA was used to analyze the data. **(I)** Energy expenditure of 6 months old control and *Myf5Cre-Pcyt2* mice during the resting (light) and active (dark) phases under thermoneutrality. **(J)** Grip strength assessment of 6 months old control and *Myf5Cre-Pcyt2* mice under thermoneutrality. Data are shown as means ± SEM. Each dot represents individual mice. *p < 0.05, **p < 0.01, ***p < 0.001, and ****p < 0.0001, n.s. not significant. Unpaired Student t-test with Welch correction was used for statistical analysis unless stated otherwise.

**Extended Data Figure 7.**
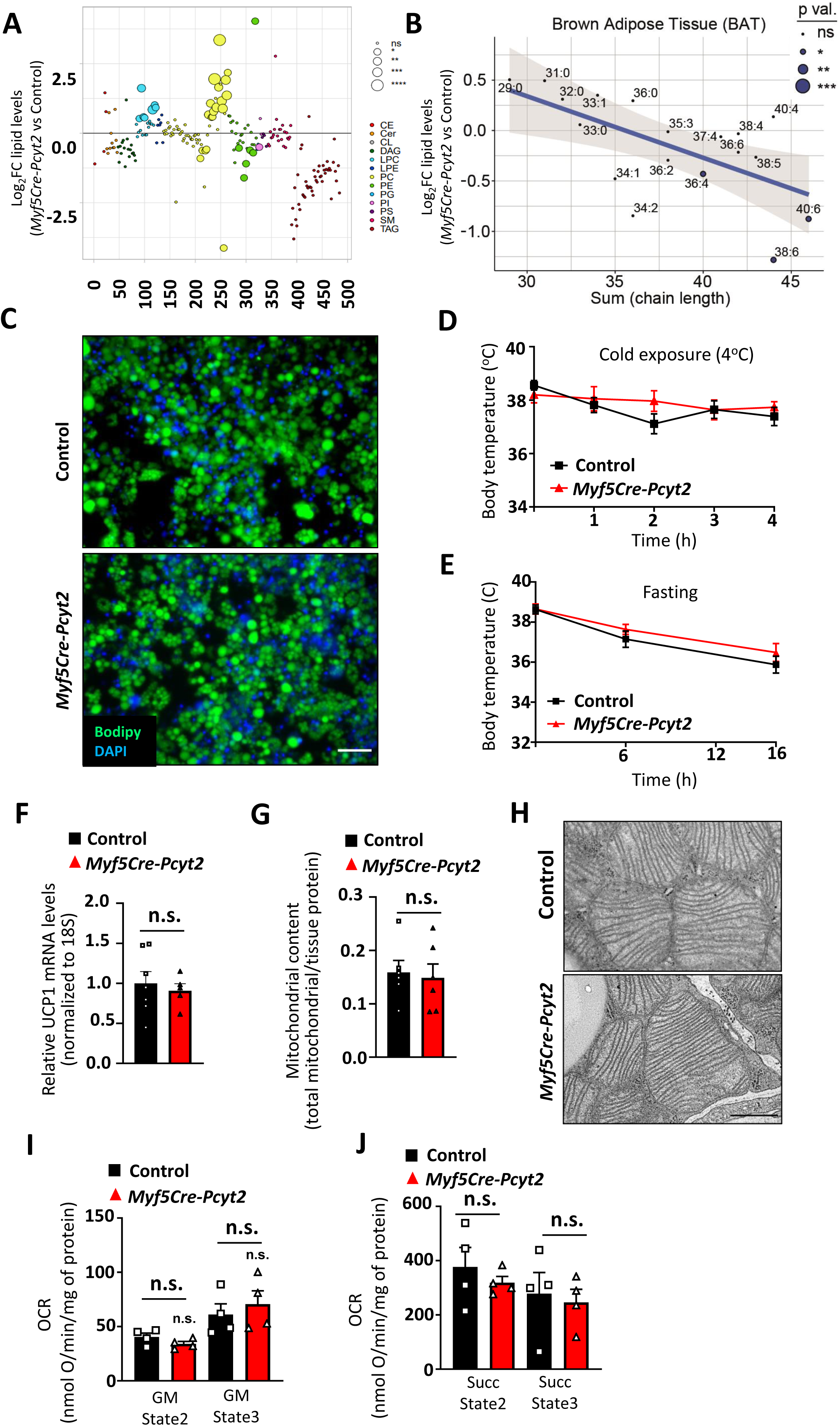
**Characterization of the brown adipose tissue from *Myf5Cre-Pcyt2* mice.** **(A-B)** Lipidomics analysis from brown fat isolated from 10-day old Myf5Cre-Pcyt2 and control mice. n=4 per group. **(C)** Brown fat differentiation in lipid free conditions from 2-day old primary pre-adipocytes isolated from control and Myf5Cre-Pcyt2 mice. Scale bar 50µm. **(D-E)** Brown fat activity as addressed by exposure of 6-month-old control and Myf5Cre-Pcyt2 mice to cold (4C) or during fasting. **(F)** Ucp1 mRNA levels in brown fat of 6-month-old control and Myf5Cre-Pcyt2 mice. **(G)** Mitochondrial content in brown adipose tissue. **(H)** BAT mitochondrial structure of 6-month-old Myf5Cre-Pcyt2 mice. Scale bar 1µm. **(I-J)** Complex I and II activities of brown fat mitochondria. Paired Student t-test was used to analyze the data. Data are shown as means ± SEM. Each dot represents individual mice. *p < 0.05, **p < 0.01, ***p < 0.001, and ****p < 0.0001, n.s. not significant (unpaired Student t-test, unless otherwise stated)

**Extended Data Figure 8.**
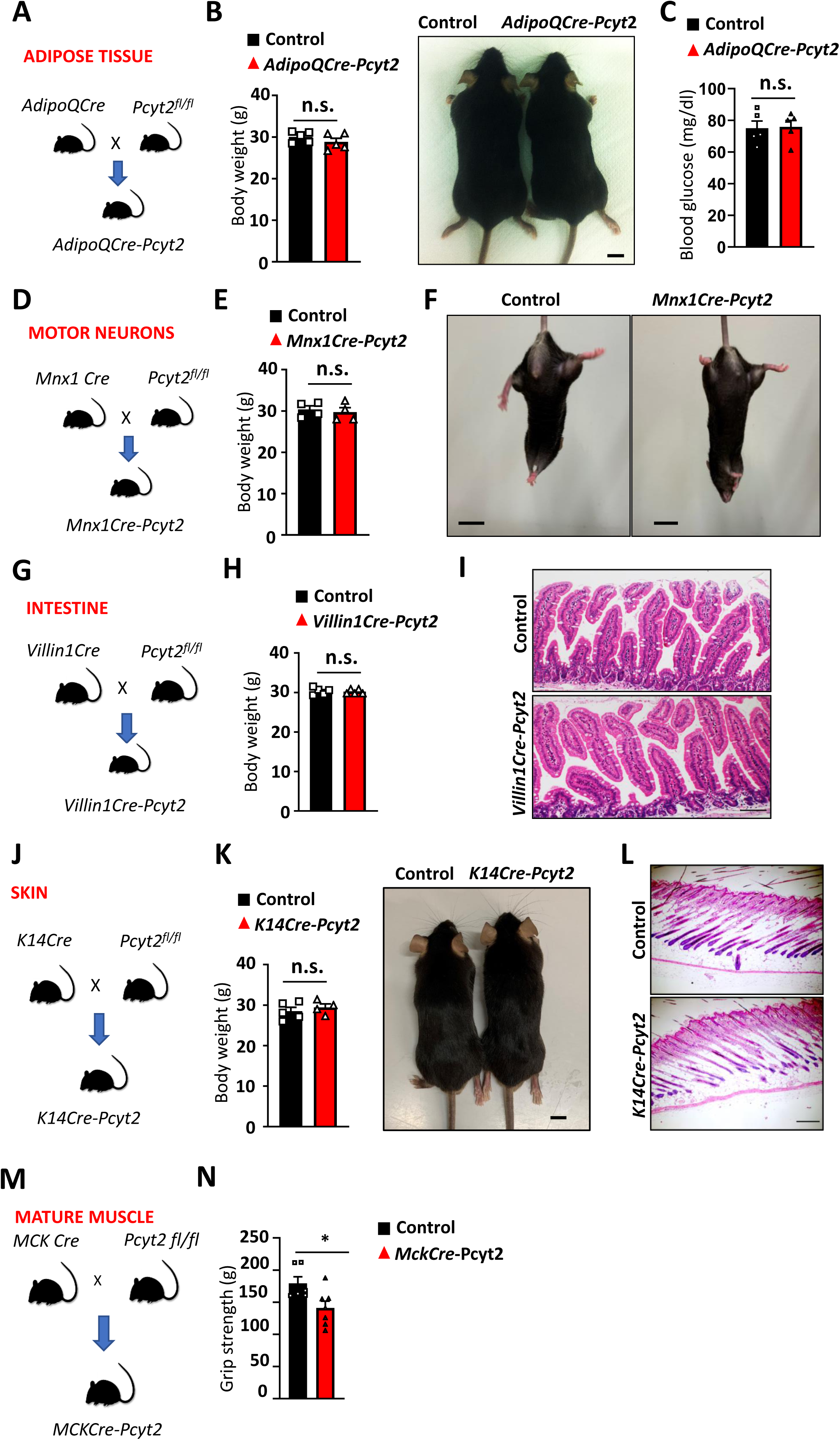
**Specific inactivation of Pcyt2 in multiple mouse tissues.** **(A)** Schematic diagram to generate adipose tissue specific *Pcyt2* deficient mice (*AdipoQCre-Pcyt2*).**(B)** Body weights and appearances of 6 months old control and *AdipoQCre-Pcyt2* mice. **(C)** Fasting blood glucose of 6 months old control and *AdipoQCre-Pcyt2* littermates fed a chow diet. **(D)** Schematic diagram of motor neuron specific *Pcyt2* deficient mice (*Mnx1Cre-Pcyt2*). **(E)** Body weights of 8 months old control and *Mnx1Cre-Pcyt2* mice. **(F)** Absence of any overt clasping behavior and appearance in 8 months old *Mnx1Cre-Pcyt2* mice. **(G)** Schematic diagram of intestine epithelium specific *Pcyt2* deficient mice (*VilinCre-Pcyt2*). **(H)** Body weights of 6 months old control and *VilinCre-Pcyt2* littermates. **(I)** Histological sections of intestine isolated from 12 months old control and *VilinCre-Pcyt2* mice. **(J)** Schematic diagram of skin epithelium *Pcyt2* deficient mice (*K14Cre-Pcyt2*). **(K)** Body weights and appearances of 6 months old control and *K14Cre-Pcyt2* mice. **(L)** Histological sections of intestine isolated from 12 months old control and *K14Cre-Pcyt2* littermates. **(M)** Schematic diagram of mature muscle specific *Pcyt2* deficient mice (*MCKCre-Pcyt2*). **(N)** Grip strength of 18 months old control and muscle specific *MckCre-Pcyt2* mice. Each dot represents individual mice, each mouse was tested in triplicates. Mean values ± SEM are displayed. *p < 0.05, **p < 0.01, ***p < 0.001, and ****p < 0.0001, n.s. not significant (unpaired Student t-test).

**Extended Data Figure 9.**
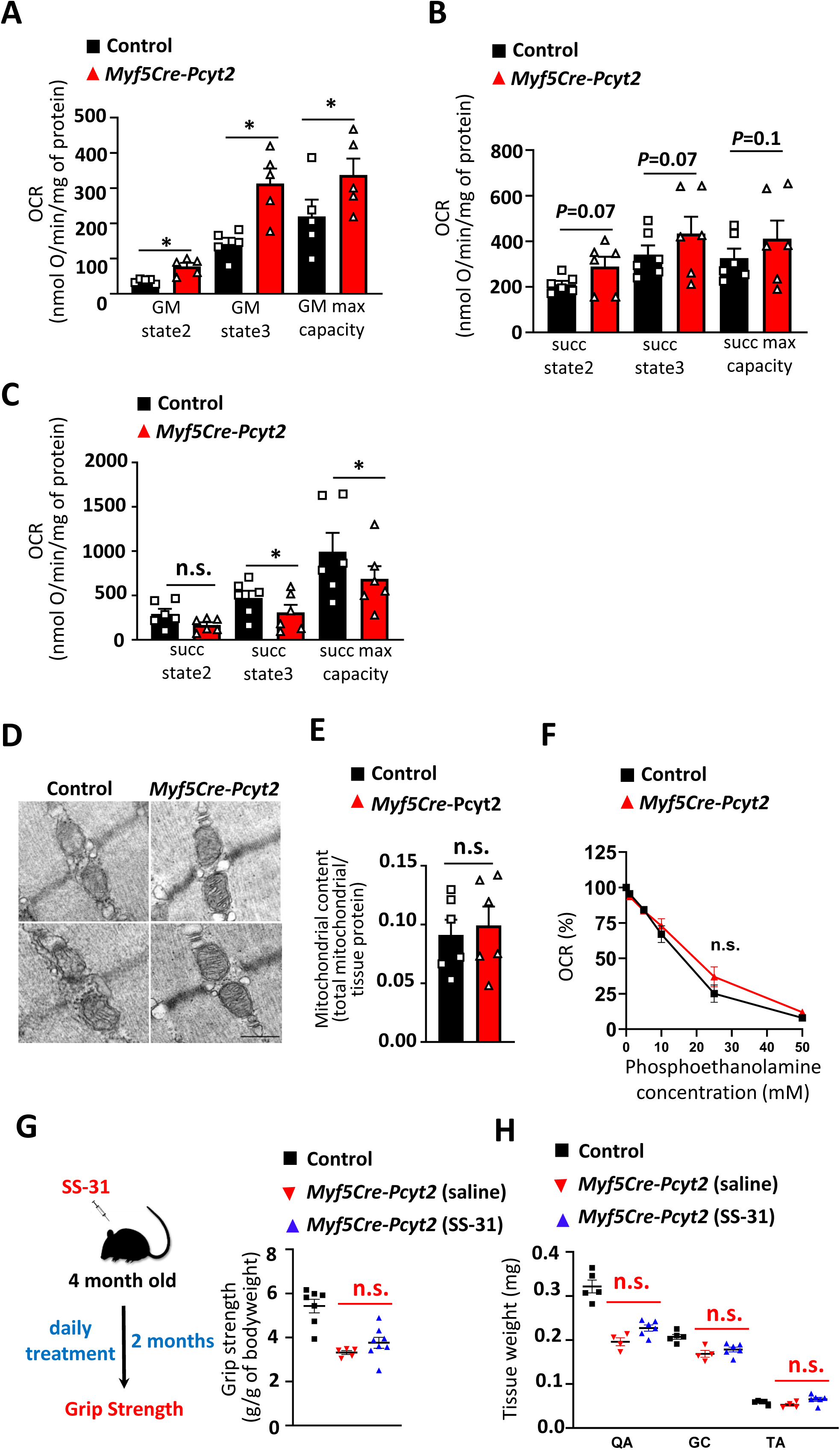
**Assessment of mitochondrial homeostasis and SS-31 treatment.** **(A-B)** Muscle mitochondrial function assessed by measurements of complex II linked activity on isolated mitochondria from 2 months and 6 months old control and *Myf5Cre-Pcyt2* mice respectively. Paired Student t-test. **(C-D)** Ultrastructure and total numbers of muscle mitochondria from 8 months -old control and *Myf5Cre-Pcyt2* mice. Scale bar 1µm. **(E)** Function of muscle mitochondria under increasing concentrations of phosphoethanolamine, as assessed by measurements of complex I linked activity on isolated mitochondria from 2 months old control and Myf5Cre-Pcyt2 mice. N=3 mice per group. **(F-G)** Grip strength and organ weight measurements of 6 months old control (vehicle) and Myf5Cre-Pcyt2 mice that have been treated with either vehicle or ss-31 compound for two months. Data are shown as means ± SEM.

**Extended Data Figure 10.**
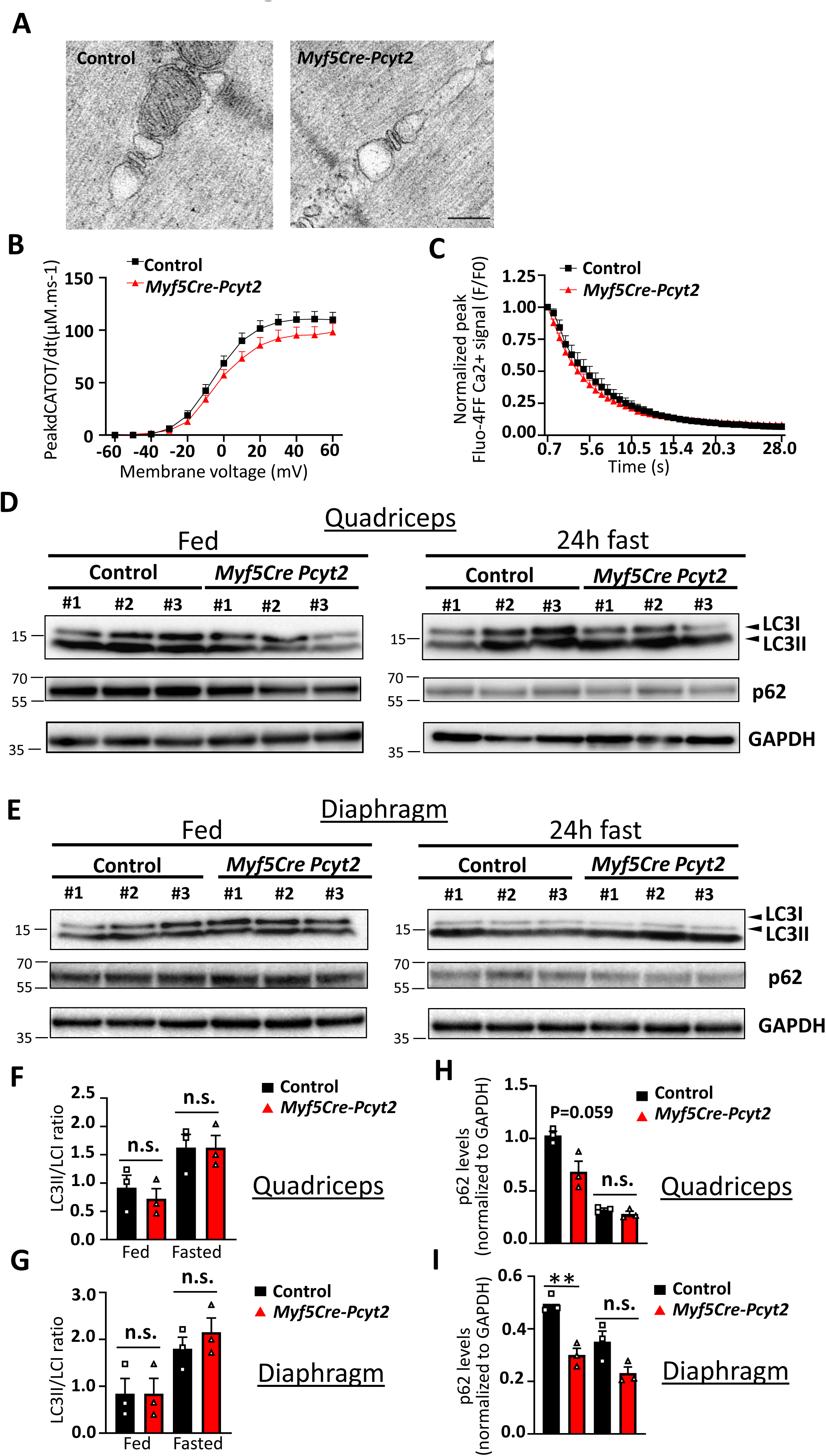
**Assessment of calcium handling, and autophagy markers in skeletal muscle.** **(A)** Representative images of SR ultrastructure in skeletal muscles from 6 months old control and *Myf5Cre-Pcyt2* mice. Scale bar 500 nm. (B) Voltage-dependence of the peak rate of sarcoplasmatic reticulum (SR) Ca^2+^ release (d[CaTot]/dt) measured from rhod-2 Ca^2+^ transients in fibers from control and *Myf5Cre-Pcyt2* mice. n=23-22 fibers isolated from n=5-6 mice per group. **(B)** Decline of voltage-activated fluo-4FF Ca^2+^ transients in control and *Myf5Cre-Pcyt2* muscle fibers in response to an exhausting voltage stimulation protocol. n=8-6 fibers isolated from n=2-3 mice per group. **(C)** LC3 I/II and p62 levels in quadriceps from 8 months old control and *Myf5Cre-Pcyt2* mice under fed and fasting (24h) conditions. N=3 mice per group. **(D)** LC3 I/II and p62 levels in diaphragm from 8 months old control and *Myf5Cre-Pcyt2* mice under fed and fasting (24h) conditions. N=3 mice per group. **(E-F)** Quantification of p62 levels under fed and fasting conditions from quadriceps and diaphragm muscle respectively. Each dot represents individual mice.

**Extended Data Figure 11.**
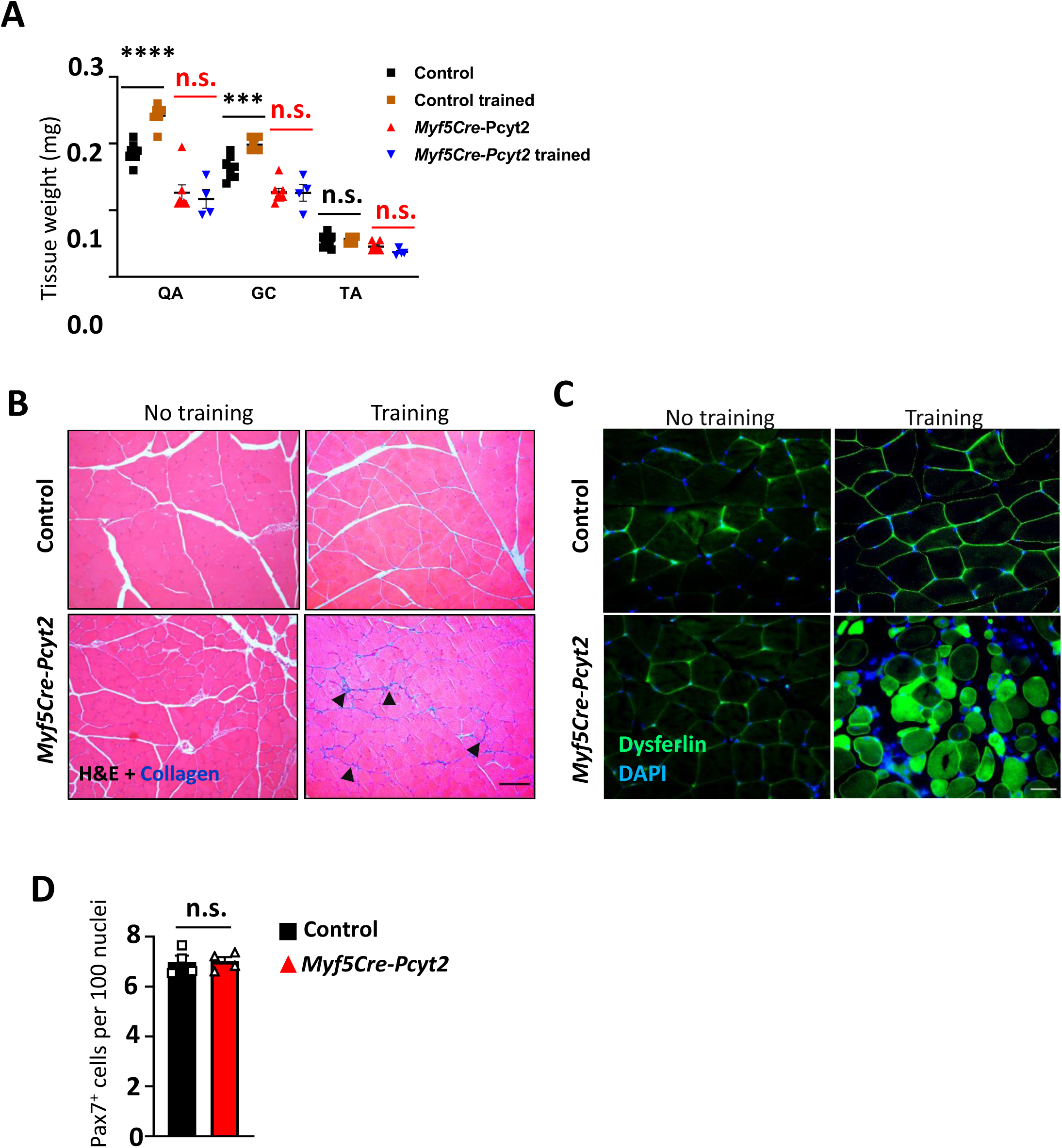
**Muscle mass, muscle damage and Pax7+ progenitor numbers after eccentric exercise in *Myf5Cre-Pcyt2* mice.** **(A)** Relative change in skeletal muscle weights from untrained and trained (3 bouts of eccentric exercise) 6 month old control and *Myf5Cre-Pcyt2* mice. QA, quadriceps; GC, gastrocnemius; TA, tibialis anterior muscles. n=4-9per group. **(B)** Tissue fibrosis (collagen staining) of quadriceps muscles isolated from 6 month old control and *Myf5Cre-Pcyt2 mice*, before (no training) and after (training) the eccentric exercise. Black arrows indicate collagen deposits. Scale bars 100µm **(C)** Dysferlin immunohistochemistry staining of the quadriceps muscles before and after eccentric exercise. DAPI was used to image nuclei. Scale bar 25µm **(D)** Number of Pax7^+^ nuclei in quadriceps muscles from 6 months old control and *Myf5Cre-Pcyt2* mice following eccentric exercise. Sections from 4 mice per group were examined, with ≥100 nuclei per section counted. Data are shown as means ± SEM. Each dot represents individual mice. *p < 0.05, **p < 0.01, ***p < 0.001, and ****p < 0.0001, n.s. not significant (unpaired Student t-test)

**Extended Data Figure 12.**
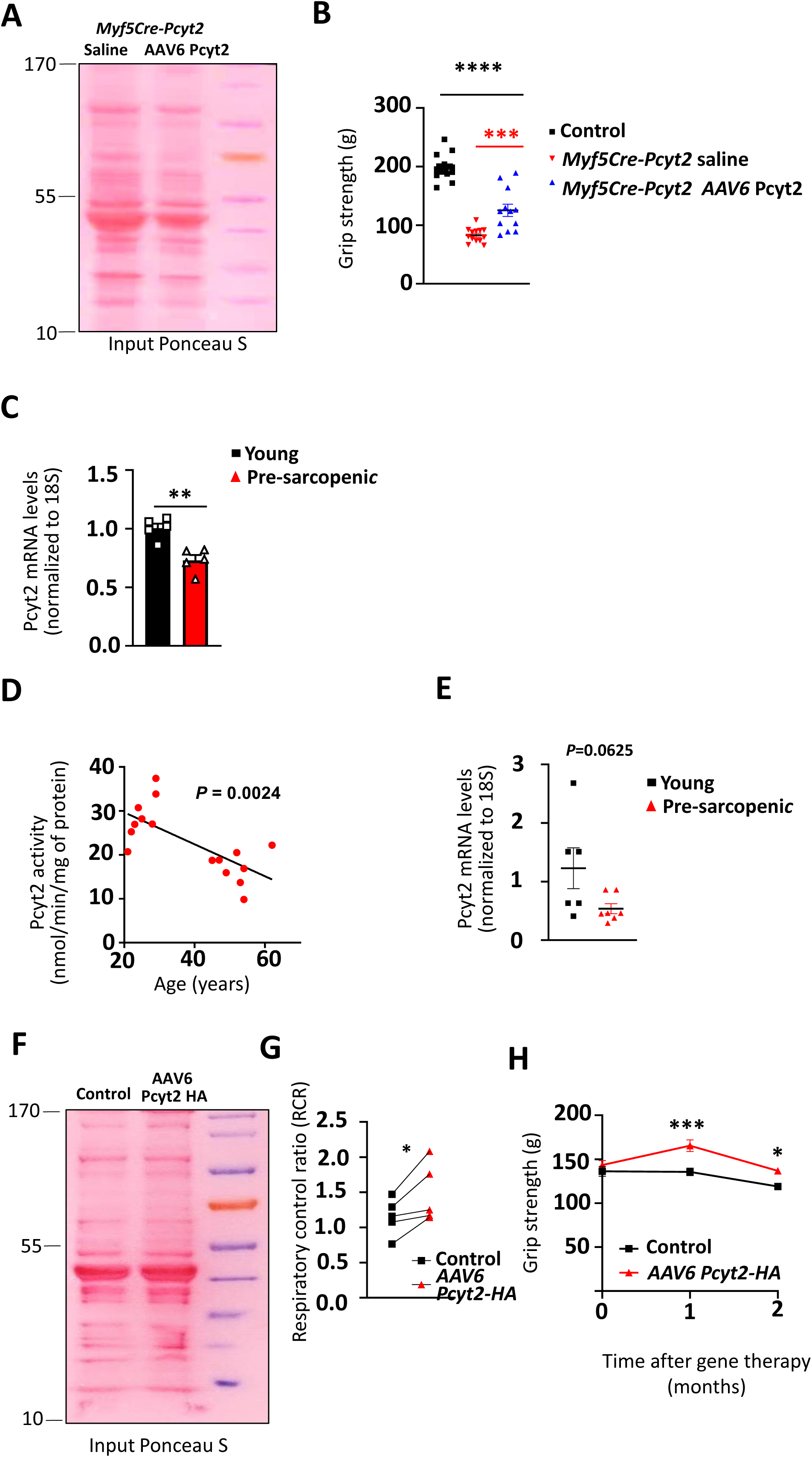
**Reduced Pcyt2 activity and levels in muscle ageing.** **(A)** Grip strength of control (saline), *Myf5Cre-Pcyt2* (saline) *and Myf5Cre-*Pcyt2 (AAV6 CK8 Pcyt2 HA) injected mice. Each dot represents individual mice. **(B)** Equal protein input is shown in the representative Ponceau S stained gel from muscle lysates of *Myf5Cre-Pcyt2* (saline) *and Myf5Cre-*Pcyt2 (AAV6 CK8 Pcyt2 HA) injected mice. **(C)** Pcyt2 mRNA levels in quadriceps muscles from young (6 month) and pre-sarcopenic (24 month old) C57B6/J mice. Each dot represents individual mouse **(D)** PCYT2 activity/age correlations and **(E)** mRNA levels in quadriceps muscle biopsied from young (20-30yr) and middle aged (45-62yr) healthy human volunteers. Each dot represents individual human. **(F)** Equal protein input is shown in the representative Ponceau S stained gel from muscle lysates of aged (24 months old) Control (saline) *and* AAV6 CK8 Pcyt2 HA injected mice 2 months after AAV6 injection. **(G)** Respiratory control ratio on muscle tissue lysates isolated from aged (24 months old) Control (saline) *and* AAV6 CK8 Pcyt2 HA injected mice 2 months after AAV6 injection.. Each dot represents individual mice. Paired Student t-test was used for statistical analysis. Dashed line indicates the average value of mitochondrial activities measured from 5 individual young (6 months old) mice. **(H)** Grip strength of 24 month old control (saline) and *AAV6-CK8-Pcyt2HA* transduced C57B6/J mice 2 months after AAV6 injection. Each dot represents individual mice. Data are shown as means ± SEM. *p < 0.05, **p < 0.01, ***p < 0.001, and ****p < 0.0001, n.s. not significant (unpaired Student t-test)

## Extended Data Video legends

### Extended Data Video 1-4

Laser induced myofiber damage assessment on myofibers isolated from 6-month-old control and Myf5Cre-Pcyt2 mice. Videos are from 2 independent isolations.

### Extended Data Video 5

Video of the last bout of eccentric exercise (day 13) of 6-month-old control and Myf5Cre-Pcyt2 mice. First, mice were adjusted to a lower speed (9m min^-1^; t=0-10s), followed by a higher speed bout (20 m min^-1^; t=10-40s).

## Methods

### CONTACT FOR REAGENT AND RESOURCE SHARING

Further information and requests for resources and reagents should be directed to and will be fulfilled by the Lead Contacts Josef M. Penninger or Domagoj Cikes.

### METHOD DETAILS

#### Studies in Humans

Patients with PCYT2 and EPT1 deficiency were identified previously ^7,9^. Their height and weight were recorded at visits to the hospital. The spinal MRI was performed at the age of 19 years. All data presented are being shared with parental and patient consent.

#### Human biopsies

All human experiments were approved by the regional ethical review board in Stockholm (2014/516-31/2 and 2010/786-31/3) and complied with the Declaration of Helsinki. Oral and written informed consent were obtained from all subjects prior to participation in the study. 8 healthy young adults (age 21-29) and 8 middle-aged (age 45-62) subjects were recruited. The subjects did not use any medications and were nonsmokers. Biopsies of the quadriceps vastus lateralis muscle were obtained under local anesthesia using the Bergström percutaneous needle biopsy technique ^82^. The biopsies were immediately frozen in isopentane, cooled in liquid nitrogen, and stored at −80°C until further analysis.

#### Studies in zebrafish

#### Generation of mutant zebrafish and analysis

Pcyt2 mutant zebrafish at F0 have been described previously ^7^. For histological examination of zebrafish models, animals were sacrificed by lethal anesthesia with buffered tricaine methanesulfonate. After gross examinations, their abdomen was opened and whole body was immersed in 4% paraformaldehyde (PFA) for 72 hr. After fixation, the animal was cut with a surgical blade into 0.5 mm pieces and embedded in paraffin blocks. 3µm sections were further processed for routine hematoxylin and eosin staining. Back muscle cross sectional areas of the same anatomical region was imaged under 10X magnification, followed by analysis (75-80 myofibers; n=4 animals per group) with ImageJ software.

#### Studies in mice

#### Mouse lines and diets

All mice were housed in the IMBA mouse colony with a 12 h light/dark cycle in a temperature-controlled environment and fed a standard chow diet. *Pcyt2* conditional mice have been described previously ^83^. In all cases, all mice described in our experiments were littermates, matched for age and sex. Tissue specific *Pcyt2* mutant mice were generated by establishing a colony of *Pcyt2^flox/flox^* mice crossed Cre transgenic mice. *Villin* Cre mice originate from Institut Curie (Sylvie Robine Lab). The following mouse lines were obtained from the Jackson Laboratory (Jackson Lab, Bar Harbor, US): *Adipoq* Cre (B6;FVB-Tg(Adipoq-cre)1Evdr/J); *Alb* Cre (B6.Cg-Speer6-ps1Tg(Alb-cre)21Mgn/J); *Mck* Cre (B6.FVB(129S4)-Tg(Ckmm-cre)5Khn/J. All animal experiments were approved by the Animal Care and Use Committee at IMBA.

#### Functional *in vivo* muscle tests

##### Grip strength

Two, 4 and 6-month old mice (control and Pcyt2 Myf5 KO mice) were subjected to grip strength tests using a grip strength meter (Bioseb, USA), following standardized operating procedures. Prior to tests, mice were single caged for two weeks, in order to avoid any littermate influence on their performance. Each mouse was tested three times, with a 15-minute inter-trial interval, and values were averaged among the three trials. Clasping index was evaluated as described previously ^84^. Each mouse was scored three times, and an average of scores was calculated.

##### Eccentric exercise

Treadmill training was performed as previously described ^85^. Briefly, single caged mice were acclimatized for treadmill exercise for three days on low speed (4m min^-1^ for 40 min per day), followed by a 7-day training on a medium speed (4m min^-1^ for 40 min plus 9m min^-1^ for 20min per day), and a 2-day stress training (20m min-1 for 20min per day) to test endurance under stress conditions. Immediately after the completion of the exercise, mice were sacrificed, and muscles were collected for histological analysis.

##### Synergic muscle ablation

For synergic ablation experiments, all surgical procedures were performed under aseptic conditions with the animals deeply anaesthetized with pentobarbital sodium (60 mg/kg i.p.). Compensatory overload of the plantaris muscle was performed unilaterally via removal of the major synergistic muscles (gastrocnemius-medialis, -lateralis and soleus), as described previously ^86^. A sham operation was systematically performed on the control hindlimb, which consisted of separating tendons of the soleus and gastrocnemius muscles from the plantaris muscle. Analgesic was administered to the animals for two days following the operation. The overload lasted for 14 days. For maintaining the activity of the animals during the overload period, moderate speed walking training was used on a treadmill (10 degree ascents, 4-5 m/min, 30 min, 6 times/week). At the end of this period, after animal sacrifice, the plantaris muscle was removed bilaterally, trimmed of excess fat and connective tissue, and wet weighed and processed for further analysis.

#### AAV-based vector delivery

For the AAV6 treatment, Pcyt2 (NM_024229.3) was C-terminally tagged with an HA-tag and cloned into the AAV6-CK8 muscle specific expression vector ^52^ using the Sal1-Kpn1 restriction sites. AAV6 viral particles were prepared as previously described ^87^. For gene therapy of Myf5Cre-Pcyt2 mutant mice, 4 day old pups were injected i.p. with 2×10^11^ vector genomes per mouse. For ageing studies 24-month-old C57B6/J mice (Jackson Labs, Bar Harbor) were single caged for two weeks for acclimatization. Grip strength was measured before AAV6 delivery using the grip strength meter. On the day of the AAV injection, mice were anesthetized with isoflurane, and injected retro-orbitally either with AAV6-Pcyt2HA (5×10^12^ vector genomes per mouse) or as a control saline. Expression of the Pcyt2-HA was determined by Western blotting and grip strength was measured one and two months after the injection.

#### Metabolic studies

Animals were fed standard chow diet and blood glucose was measured at fed and fasted state (16h fasting). Standard chow diet was purchased from SSNIFF (V 1184-300; 10mm pellets autoclavable; 49% kcal carbohydrates, 36% kcal protein and 15% kcal fat). Measurements were done at the same time of the day, by collecting blood samples after tail snipping and applying the blood samples to Onetouch Verio strips (LifeScan, GmbH). Food consumption was measured on single cage housed animals over a period of two weeks, after acclimatization single caging for one week. For calorimetry, measurements were performed at room temperature (21C-23C) on a 12/12 h light/dark cycle in a PhenoMaster System (TSEsystems, Bad Homburg, Germany) using an open circuit calorimetry system ^88^. Mice (4-7 months old) were housed individually, trained on drinking nozzles for 7 days and allowed to adapt to the PhenoMaster chamber for 2 days. Food and water were provided ad libitum. Parameters of indirect calorimetry and activity were determined for 5 consecutive days. Body weights were recorded at the beginning and end of the experiments and average values were plotted against energy expenditure and activity ^89^. To address brown fat activity, 6-month-old mice were housed at 4°C and body temperature was measured during the fed period using a thermometer for small animals (Thermometer TK 98802; Bioseb). Temperature was recorded every hour over a 4 h period. After a 2 day recovery period at room temperature, the same mice were fasted and the body temperatures determined at 6 and 16 hours of fasting. To determine the autophagic response, animals were fasted for 24 hours.

#### Primary cell culture conditions

##### Myoblasts

Primary myoblasts were isolated as previously described ^90^. Briefly, after collagenase (type 2; 500U per mL) and liberase digestion (2.5U/mL) for 1.5hr on 37°C, the tissue slurry was seeded directly on Matrigel coated dishes (45 μg/cm2) in DMEM/F12 containing 20%FCS (Sigma-Aldrich) and 10% horse serum (Sigma-Aldrich) and penicillin– streptomycin (Life Technologies, final conc., 50 U/mL of penicillin). Removal of fibroblasts was achieved by serial (2-4) passaging and pre-plating the cell suspension on collagen Type 1 coated dishes for 1 hr prior transfer to Matrigel coated dishes (4.5 μg/cm2). Differentiation into myotubes was done by culturing myoblasts in serum/lipid free DMEM/F12 with Insulin-Transferrin-Selenium supplement (ThermoFisher, GmbH) for either 2 days to examine the levels of early differentiation and fusion markers, or for 5 days to analyze the fusion efficiency.

To evaluate the influence of masking the outer membrane phosphatydilethanolamine, primary myoblast differentiation was done in myoblast differentiation media desrcibed above with addition of either vehicle (PBS) or SA-Ro phosphatydilethanolamine binding probe (2.5µg/mL final concentration). The media containing fresh SA-Ro probe was exchanged every two days, and the fusion efficiency as evaluated on day 5.

##### Brown adipocytes

Primary brown adipocyte differentiation was done as previously described ^91^. Briefly, after harvesting from 2-day old pups, brown fat tissue was digested in DMEM collagenase solution (type 2; 500U per mL) for 1.5hr. Cells were grown in DMEM with 10% FCS and penicillin–streptomycin (Life Technologies, final conc., 50 U/mL of penicillin). Differentiation into mature brown fat adipocytes was achieved by adding an adipogenic cocktail (0.5 μg/mL insulin, 5 μM dexamethasone, 1 μM rosiglitazone, and 0.5 mM IBMX) in serum free media for 3 days, following maintenance in serum free media supplemented only with insulin for another 4 days. Lipid droplets were visualized with Bodipy 493/503 solution (ThermoFisher).

#### Lipidomics

Quadriceps muscles and brown adipose tissues were isolated from 10-day old pups and snap frozen in liquid nitrogen. Muscle tissue was homogenized using a Precellys 24 tissue homogenizer (Precellys CK14 lysing kit, Bertin). Per mg tissue, 3µL of methanol were added. 20 µL of the homogenized tissue sample was transferred into a glass vial, into which 10 µL internal standard solution (SPLASH® Lipidomix®, Avanti Polar Lipids) and 120 µL methanol were added. After vortexing, 500 µL Methyl-tert-butyl ether (MTBE) were added and incubated in a shaker for 10 min at room temperature. Phase separation was performed by adding 145 µL MS-grade water. After 10 min of incubation at room temperature, samples were centrifuged at 1000 x*g* for 10min. An aliquot of 450 µL of the upper phase (organic) was collected and dried in a vacuum concentrator. The samples were reconstituted in 200 µL methanol and used for LC-MS analysis. The LC-MS analysis was performed using a Vanquish UHPLC system (Thermo Fisher Scientific) combined with an Orbitrap Fusion™ Lumos™ Tribrid™ mass spectrometer (Thermo Fisher Scientific). Lipid separation was performed by reversed phase chromatography employing an Accucore C18, 2.6 µm, 150 × 2 mm (Thermo Fisher Scientific) analytical column at a column temperature of 35°C. As mobile phase A we used an acetonitrile/water (50/50, v/v) solution containing 10 mM ammonium formate and 0.1 % formic acid. Mobile phase B consisted of acetonitrile/isopropanol/water (10/88/2, v/v/v) containing 10 mM ammonium formate and 0.1% formic acid. The flow rate was set to 400 µL/min. A gradient of mobile phase B was applied to ensure optimal separation of the analysed lipid species. The mass spectrometer was operated in ESI-positive and -negative mode, capillary voltage 3500 V (positive) and 3000 V (negative), vaporize temperature 320°C, ion transfer tube temperature 285°C, sheath gas 60 arbitrary units, aux gas 20 arbitrary units and sweep gas 1 arbitrary unit. The Orbitrap MS scan mode at 120000 mass resolution was employed for lipid detection. The scan range was set to 250-1200 m/z for both positive and negative ionization mode. The AGC target was set to 2.0e5 and the intensity threshold to 5.0e3. Data analysis was performed using the TraceFinder software (ThermoFisher Scientific). The lipidomics results from five biological replicates per group were analyzed as reported previously ^92^. Briefly, the amount of each lipid was normalized to the sum of concentrations for all lipid species, measured in a single biological replicate. Values were next averaged over the five biological replicates for control and Myf5Cre-Pcyt2 samples, log2 transformed, and compared between the groups.

To determine total phosphatydilethanolamine levels in skeletal muscles, quadriceps muscles from either 6 month old control, *Myf5Cre-Pcyt2* mice (saline), *Myf5Cre-Pcyt2* mice (AAV6-*Pcyt2*) or from young (6 month old), old (24 month old, saline), old (24 month old, AAV6-*Pcyt2*) were surgically removed, snap frozen in liquid nitrogen, and grinded to powder on dry ice. The weight of each sample was than measured using a microscale. Total phosphatydilethanolamine levels were than analyzed using Phosphatidylethanolamine Assay Kit (ab241005, Abcam, UK) according to the manufacturers’ recommendations. Final levels were determined after normalization to the sample weight.

To determine significant changes un-paired t-test was used and represented as ns - non significant, *\*P* <0.05, *\*\*P* <0.01, *\*\*\*P* <0.001, *\*\*\*\*P* <0.0001.

#### Respiration and catalase activity measurements

Mitochondrial respiratory parameters were measured with high-resolution respirometry (Oxygraph-2k, Oroboros Instruments, Innsbruck, Austria). Briefly, either tissue lysates or isolated mitochondria (0.03 mg/mL) were incubated in a buffer containing 80 mM KCl, 5 mM KH_2_PO_4_mM, 50 mM Mops, 1mM ethylene glycol-bis(2-aminoethylether)-N,N,N′,N′-tetraacetic acid, and 1 mg/mL fatty acid-free bovine serum albumin (pH 7.4, 37 °C)1. Respiration was initiated either by the addition of 5 mM glutamate and 5 mM malate (complex I), 10 mM pyruvate (complex I), or 10 mM succinate in the presence of 1.4 µM rotenone (complex II). Transition to State-3 respiration was induced by the addition of 1 mM adenosine diphosphate. Maximum electron transfer system capacity was measured by titration of carbonyl cyanide-4-(trifluoromethoxy) phenylhydrazone in steps of 0.1 µM. Addition of exogenous cytochrome c (2.5 µM) was used to estimate the permeability of the outer mitochondrial membrane as a quality control. Oxygen consumption rates were obtained by calculating the negative time derivative of the measured oxygen concentration and normalized to total protein amount of the mitochondrial suspension or the tissue lysate. Catalase activity was determined from isolated quadriceps of 6-month-old mice using a catalase activity assay according to the manufacturer recommendations (#700910, Cayman Chemical).

#### Isolation and imaging of Giant Plasma Membrane Vesicles (GPMVs)

GPMVs were prepared as previously described ^93^. Briefly, myoblasts were seeded on a 60 mm petri dish until ∼ 70% confluency. Before GPMV formation, they were washed twice with GPMV buffer (150 mM NaCl, 10 mM Hepes, 2 mM CaCl_2_, pH 7.4) and finally 2 ml of GPMV buffer was added to the cells together with 25 mM PFA and 10 mM DTT (final concentrations). After incubation for 2hr at 37°C, GPMVs were collected from the supernatant. For GMPV preparation from myofibers, the Extensor digitorum longus (EDL) muscle was digested in collagenase supplemented medium (type1, 2mg/mL) for 2.5hr at 37°C, followed by trituration and single myofiber isolation as described above. Myofibers were then gently washed twice with FCS free DMEM, followed by a brief 1min wash with GPMV buffer containing 25 mM PFA, to prevent myofiber hypercontraction. Finally, GPMV buffer with 25 mM PFA and 10 mM DTT was added to the myofibers. After incubation for 2hr at 37°C, GPMVs were collected from the supernatant. GMPVs were labelled with the polarity sensitive membrane probe NR12S (a kind gift from A. Klymchenko) at 0.1µg/ml final probe concentration in phosphate buffered saline (PBS) for 5 minutes and then immediately imaged at room temperature. Spectral imaging was performed on a Zeiss LSM 780 confocal microscope equipped with a 32-channel GaAsP detector array. Laser light at 488 nm was used for fluorescence excitation of NR12S. The lambda detection range was set between 490 - 691 nm. Images were saved in lsm file format and then analyzed by using a custom plug-in compatible with Fiji/ImageJ3 ^94^. Glass coverslips (thickness of 0.17 mm) were used to measure generalized polarization which reflects membrane lipid packing/order using the following formula where I_560_ and I_650_ are the fluorescence intensities at 560 nm and 650 nm respectively:

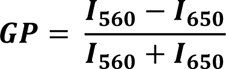

#### Brillouin Microscopy of myofibers

Brillouin Light Scattering Microscopy (BLSM) was performed using an inverted confocal sample-scanning microscope with a Brillouin imaging-spectrometer as described by us previously ^45,95^. Briefly the setup employed a 532nm single-frequency probing laser and is based on a 2-stage cross dispersion Virtual Imaged Phase Array (VIPA) with intermediate Fourier and image plane filtering, a cooled EM-CCD camera (Hamamatsu ImagEMII) for detection ^96,97^, and can achieve a spectral finesse >85. For more scattering samples, we also employed a heated Iodine absorption cell in the detection path tuned to the laser wavelength to reduce the elastic scattering signal ^98^. To generate spatial maps, samples were scanned with a 3-axis piezo electric stage (Physik Instrumente). Imaging was performed through 1.3 Numerical Aperture (NA) Si immersion-oil objective (Olympus) and confocallity was assured via a physical pinhole of ∼1 Airy Unit (AU) before coupling the light into the spectrometer. Widefield transmitted light images were used to determine the scanning area for each sample. Several cross-sectional scans were performed for each myofiber at positions separated by a 1µm. Acquisition was controlled by a custom Labview based script developed by the company THATec. The acquisition (dwell) time per voxel was 100ms and the power measured at the sample was 2-3mW. Each measured spectra was de-convolved with the complete system spectral response as determined for the attenuated elastic scattering peak measured prior to each scan in the same sample. Prior and subsequent to each imaging session the spectra of water and ethanol were measured on a separate imaging arm and used for calibrating the spectral projection based on paraxial theory ^99^, adapted to a dual VIPA setup. Each spectrum was then analyzed in Matlab (Mathworks) using custom developed scripts employing Spectral Phasor Analysis ^100^ Results were confirmed to be in agreement with ones obtained from conventional non-linear least-squares fitting of deconvolved spectra consisting of the Stokes and Anti-Stokes Brillouin peaks using Lorentzian functions, however yielding lower uncertainties on average. The extracted Brillouin peak frequency, which scales with the local elastic storage modulus, is taken to be indicative of the local stiffness. All measurements were performed at 37°C and 5% CO2.

#### Atomic force microscopy

Force spectroscopy by means of Atomic force microscopy was done on individual myofibers after isolation from EDL muscles. Fibers were either cultured on matrigel coated dishes (QI mode) or probed immediately after isolation (Force spectroscopy). A JPK (Bruker) Nanowizard4 AFM atomic force microscope was used for the AFM experiments. This system implements the Quantitative Imaging (QI) mode, where a force-curve is acquired at each pixel location of the imaged area, providing stiffness data together with topography. The QP-BioAC cantilevers from Nanosensors (0.06N/m, less than 10 nm nominal tip radius) were used because of their ability to work in QI mode with biological samples. The approach and retract speeds were kept constant at 52microns/s. The model used to obtain the Young’s modulus from the acquired data was Hertz/Sneddon, as implemented in the JPK software. The paraboloid model was chosen as the most suitable for the sharp tips of the QP-BioAC cantilevers. For the fits forces up to 60pN were considered, where indentation depths remained below 500nm. Each of the fibers retrieved ≥ 4000 values (5µm X 5µm sampled in 64×64 data pixels). Matlab’s Randsample function was used to uniformly sample 500 data points.

For the force curves acquired for the qualitative comparison, forces up to 600pN were applied to a single location each, using same cantilevers and approach speeds. Samples were kept at a 37°C during the experiment.

#### Laser-induced damage of myofiber cell membranes

Myofibers were freshly isolated from EDL muscle of 4-month old mice by digestion in collagenase supplemented media (type1, 2mg/ml, Sigma) for 2 hr at 37°C. After digestion, individual fibers were transferred onto a 4-well chamber slide (Nunc Lab-Tek, Merck, GmbH) containing HBSS with 2.5μM FM1-43 dye (Molecular Probes, Invitrogen, ThermoFisher Scientific, GmbH). Laser-induced cell membrane damage was performed as previously described ^48^. Briefly, a 5 × 5 pixel area of the plasma membrane was exposed to a Laser at 20% of maximum power (Enterprise, 80 mW, 351/364 nm) for 5 s using a Zeiss-LSM 510 confocal microscope equipped with a ×63 water immersion lens (N.A. 1.3). Following the laser damage, distribution of the FM1-43 dye was imaged using high speed video captures. Throughout the experiment, cells were kept at a 37°C in a 5% CO_2_ chamber.

#### Ca2+ dynamics under voltage-clamp protocol

The flexor digitorum brevis (FDB) and interosseous (IO) muscles were freshly isolated from mice and incubated with collagenase (Sigma type 1) for 1 hr at 37 °C in a Tyrode solution. Single myofibers were isolated by gentle mechanical trituration of the collagenase-treated muscles within the experimental chamber. Fibers were voltage-clamped using the silicone-voltage-clamp technique as described (Jacquemond, 1997; Lefebvre et al., 2014) with the voltage-clamp pipette filled with a solution containing (in mM) 140 K-glutamate, 5 Na2-ATP, 5 Na2-phosphocreatine, 5.5 MgCl2, 5 glucose, 5 HEPES and either 15 EGTA, 6 CaCl2, and 0.1 rhod-2 or 0.1 fluo-4FF. The extracellular solution contained (in mM) 140 TEA-methane-sulfonate, 2.5 CaCl2, 2 MgCl2, 1 4-aminopyridine, 10 HEPES and 0.002 tetrodotoxin. For fluo-4FF fluorescence measurements, the extracellular solution also contained 0.05 mM N-benzyl-p-toluene sulfonamide (BTS) to block contraction. All solutions were at pH 7.2. Voltage-clamp and membrane current measurements were done with an RK-400 patch-clamp amplifier (Bio-Logic, Claix, France) in whole-cell voltage-clamp configuration. Command voltage pulse generation was achieved with an analog-digital converter (Digidata 1440A, Axon Instruments, Foster City, CA) controlled by pClamp 9 software (Axon Instruments). Analog compensation was adjusted to decrease the effective series resistance. Holding voltage was always set to −80 mV. Following insertion of the micropipette extremity into the muscle fiber, a 30 min-long period of equilibration was allowed before taking measurements. Rhod-2 and fluo-4-FF fluorescence were detected with a Zeiss LSM 5 Exciter confocal microscope equipped with a 63× oil immersion objective (numerical aperture 1.4). For detection of rhod-2 and fluo-4 FF fluorescence, excitation was from the 543 nm line of a HeNe laser and from the 488 nm line of an Argon laser, respectively, and fluorescence was collected above 560 nm and above 505 nm, respectively. Both rhod-2 and fluo-4 FF voltage-activated fluorescence changes were imaged using the line-scan mode (x,t) of the system and expressed as F/F0 where F0 is the baseline fluorescence. Rhod-2 Ca2+ transients were triggered by 0.5 s-long depolarizing pulses of increasing amplitude. From these, the rate of SR calcium release (d[CaTot]/dt) was calculated as described (Kutchukian et al., 2016; Lefebvre et al., 2014). Fluo-4-FF was used under non-EGTA buffering intracellular conditions to assess the resistance of the fibers to a fatigue protocol. For this, fibers were stimulated by consecutive trains of thirty 5 ms-long pulses from −80 mV to +60 mV delivered at 100 Hz: 40 trains were applied, separated by a 0.7 s interval.

#### Immunoblotting and Elisa measurements

##### Western blotting

For immunoblotting analyses, tissues were snap frozen and homogenized in RIPA buffer (Sigma; R0278) containing Halt protease and phosphatase inhibitor cocktail (Thermo Scientific; 78440) immediately after isolation from sacrificed animals. Protein levels were determined using the Bradford assay kit (Pierce, GmbH) and lysates containing equal amounts of protein were subjected to SDS-PAGE, further transferred to nitrocellulose membranes. Western blotting was carried out using standard protocols. Blots were blocked for 1 hour with 5% BSA (Sigma Aldrich; #820204) in TBST (1× TBS and 0.1% Tween-20) and were then incubated overnight at 4°C with primary antibodies diluted in 5% BSA in TBST (1:250 dilution). Blots were washed three times in TBST for 15 min, then incubated with HRP-conjugated secondary antibodies diluted in 5% BSA in TBST (1:5000 dilution) for 45 min at room temperature, washed three times in TBST for 15 min and visualized using enhanced chemiluminescence (ECL Plus, Pierce, 1896327). The following primary antibodies were used: anti-phospho-JNK (#4668 CST, DE, 1:150), anti-total JNK (#9255 CST, DE, 1:150), anti-FoxO1 (#2880 CST, DE, 1:150), anti-LC3 (#ab51520 Abcam, UK, 1:150), anti-pRPS6 (#4858, CST CST, DE, 1:1000). anti-RPS6 #2217 CST, DE, 1:1000), anti-peIF4A1(# PA5-105329, Thermo Fisher, DE 1:500), anti-eIF4A(#2013, CST, DE, 1:500), anti-LC3I/II (#2775 CST, DE, 1:1000), anti p62(# 600-401-HB8, ThermoFisher DE, 1:1000), anti-GAPDH (#8884 CST, DE, 1:1000). Secondary antibodies were anti-rabbit IgG HRP (CST, NE,#7074)

##### Carbonylated proteins and myofiber mtROS detection

To detect carbonylated proteins, snap frozen tissues were lysed in RIPA buffer and further processed using the OxyBlot Protein Oxidation Detection Kit (EMD Milipore, GmbH) following the manufacturer’s protocol. The carbonylated protein signal was normalized to total protein levels determined by Ponceau S staining using densitometry measurements with ImageJ software.

To detect myofiber mtROS levels, myofibers were freshly isolated from EDL muscle of 6-month old mice as described above. Immediately after the isolation, mtROS MitoSOX probe (#M36008, Thermofisher Scientific, DE) and Hoechst stain were added to live myofibers in HBSS/Ca^2+^/Mg^2+^ according to the manufacturers’ instructions. Myofibers were than imaged, and fluorescence intensity of both probes was calculated using ImageJ software. Final levels of mtROS were determined after normalizing the intensity of MitoSOX to the intensity of Hoechst of each individual myofiber.

##### Immunoprecipitation

For immunoprecipitation experiments, mice were sacrificed by cervical dislocation after completion of the analysis. Muscles were surgically removed and snap frozen in liquid nitrogen. The tissues were homogenized in RIPA buffer containing protease and phosphatase inhibitors. Equal amount of protein, as determined by Bradford assay kit (Pierce, GmbH), was used for anti-HA immunoprecipitation following manufacturer recommendations (#26180, Thermo Fisher Scientific, Gmbh). After immunoprecipitations, samples were subjected to SDS-PAGE, and further transferred to nitrocellulose membranes, incubated with primary anti-Pcyt2 antibody (HPA023033, Atlas Antibodies, Sweden) Followed by and a HRP-labelled secondary Anti-rabbit antibody (#7074, CST, NE, #7074).

##### ELISA

Tissue lysates in RIPA buffer containing protease and phosphatase inhibitors as above were used for ELISA measurements of IL-1 beta (#RAB0275, Sigma Aldrich, GmbH) and IL-6 (#RAB0309, Sigma Aldrich, GmbH) following manufacturer’s recommendations. Absorbance values were measured using the Tecan microplate reader and data were processed using the Magellan software. Concentrations were determined based on standard curves and normalized to total protein concentrations of the corresponding sample determined by Bradford assay.

Blood muscle creatine kinase levels were determined using Creatine Kinase Activity Assay Kit (ab155901, Abcam, UK) according to the manufracturers’ instructions.

#### RNA tissue extraction and Quantitative real-time PCR (RT-qPCR)

Muscles or muscle biopsies were freshly isolated and flash frozen in liquid nitrogen. The mRNA from tissues or primary myoblast cultures was isolated using the RNeasy Lipid Tissue Mini Kit (Qiagen, GmbH). RNA concentration was estimated by measuring the absorbance at 260 nm using Nanodrop (Thermofisher, GmbH). cDNA synthesis was performed using the iScript Advanced cDNA Synthesis Kit for RT-qPCR (Bio-Rad, GmbH) following manufacturer’s recommendations. cDNA was diluted in DNase-free water (1:10) before quantification by real-time PCR. mRNA transcript levels were measured in triplicate using the the CFX384 Real-Time PCR Detection System (BioRad) using the SybrGreen qPCR Master Mix (BioRad. GmBH). . Detection of the PCR products was attained with SYBR Green (Bio-Rad, GmbH). After completion of each run, melting curve analyses were performed to check for unspecific products, and representative samples run on agarose gels to ensure the specificity of amplification. Gene expression was normalized to the expression level of 18S ribosomal rRNA as the reference gene. For normalization, threshold cycles (Ct-values) were normalized to 18 ribosomal rRNA within each sample to obtain sample-specific ΔCt values (= Ct gene of interest – Ct housekeeping gene). 2^-ΔΔCt^ values were calculated to obtain fold expression levels. Primers sequences for myoblast cell culture experiments were published previously^101^. The additional primer sequences are as follows:

##### 18SrRNA

FW 5’- GGCCGTTCTTAGTTGGTGGAGCG -3’

RV 5’- CTGAACGCCACTTGTCCCTC - 3’

##### Pcyt2

FW 5’- CCGGTACAAGGGCAAGAACT -3’

RV 5’- TCTTCTCTTGGGCTCCTGGT - 3’

##### Atrogin1

FW 5’- GCAAACACTGCCACATTCTCTC -3’

RV 5’- CTTGAGGGGAAAGTGAGACG - 3’

##### MuRF1

FW 5’- ACCTGCTGGTGGAAAACATC -3’

RV 5’- ACCTGCTGGTGGAAAACATC - 3’

##### FbxO31

FW 5’- GTATGGCGTTTGTGAGAACC -3’

RV 5’- AGCCCCAAAATGTGTCTGTA - 3’

##### 18SrRNA (Human)

FW 5’- GGCCCTGTAATTGGAATGAGTC -3’

RV 5’- CCAAGATCCAACTACGAGCTT - 3’

##### PCYT2 (Human)

FW 5’- GGTGCGATGGCTGCTATGA -3’

RV 5’- CACCTCGTCCACCCATTTGA - 3’

#### Histology, immunohistochemistry, and electron microscopy

##### Histology

Muscles were isolated from sacrificed animals and fixed in 4%PFA for 72 hr. Tissues were embedded in paraffin blocks and 2µm sections were H&E stained for histopathological examination. To address the effect of the 14-day eccentric exercise regime, animals were immediately sacrificed upon completion of the last exercise bout, muscles isolated and fixed in 4% PFA, and further processed for H&E and toluidine blue staining. For determination of central nuclei quadriceps cross sections (n=4 per group) were H&E stained and 200-400 nuclei were counted. For extracellular collagen detection, 3µm paraffin muscle sections were stained with Massone Trichrome staining kit according to manufacturer’s recommendations (Abcam, UK, ab150686).

##### Immunocytochemistry

For immunocytochemistry, quadriceps muscles were harvested and fixed in PFA (4%) for 72h and cryoprotected by further immersing in 30% sucrose solution for another 72 hr. After embedding in OCT, sections were cut and stained using standard immunohistochemistry using the following antibodies: anti-Dystrophin (#ab15277, Abcam, UK1:150), anti-Pax7 (P3U1, DSHB, Iowa, US 1:150), antiDysferlin (#ab124684, Abcam, UK); anti-CD3 (#ab49943, Abcam, UK 1:3000), anti-CD4 (#14-9766-82, eBioscience, ThermoFisher, GmbH, 1:100), anti-CD8 (#14-0808-82, ThermoFisher, GmbH, 1:100), Ly6G (#127602, BioLegend, US, 1:500), anti-ClC3 (#9661S, Cell Signaling, DE, 1:100), anti-B220 (#550286, BD Biosciences, US, 1:50), and anti-F4/80 (#MCA497G, BioRad, GmbH, 1:100). For F-actin staining, longitudinal muscle sections were stained for 15 minutes with Phalloidin– Atto 565 fluorescent label (Sigma Aldrich, GmBH, 1:10000) and imaged using Zeiss LSM 780.

For analysis of mean fiber area, quadriceps cross-sectional area and mean muscular fiber number, dystrophin stained muscle sections were drawn, segmented, and analyzed using Fiji software.

For immunocytochemistry analysis of myofiber types, quadriceps muscles were harvested and frozen by immersing the unfixed quadriceps muscle in OCT and slowly freezing the block in pre-cooled isopentane solution in liquid nitrogen. Sections were cut and stained using standard immunohistochemistry using the following antibodies: BA-D5, SC-71 and BF-F3 (DSHB, Iowa, US 1:150) and counterstained with DAPI.

For analysis of glycolytic versus oxidative fiber type, unfixed muscle (quadriceps) was cut and stained as reported^102^. A large area was imaged under 10X magnification, and the analysis for the number of glycolytic and xidative fibers was done manually.

##### BrdU and EdU labelling

Two day old pups from control and Myf5-Pcyt2 pups were injected with BrdU (150 mg/kg) and sacrificed 4 hours after injection. Quadriceps muscles were surgically removed and processed for immunofluorescence as described above. Percentage of BrdU^+^ nuclei was determined after normalizing the BrdU^+^ nuclei to DAPI stained nuclei.

For EdU labeling, primary myoblasts were labelled for 90 minutes and processed with EdU Click-iT™ EdU Cell Proliferation Kit (C10337, ThermoFisher, GmbH). Percentage of EdU^+^ nuclei was determined after normalizing the EdU^+^ nuclei to DAPI stained nuclei.

##### Electron microcopy

For electron microcopy analysis, 6 month old animals were sacrificed, muscles were surgically excised, trimmed to small pieces and immediately immersed in cold 2.5% glutaraldehyde. Muscles were processed for electron microsopy as described previously

^103^. Briefly, tissue was cut into small pieces and fixed in 2.5% glutaraldehyde in 0.1 mol/l sodium phosphate buffer, pH 7.4. for 1 hour. Subsequently samples were rinsed with the same buffer and post-fixed in 2% osmium tetroxide in 0.1 mol/l sodium phosphate buffer on ice for 40 min. After 3 rinsing steps in ddH2O, the samples were dehydrated in a graded series of acetone on ice and embedded in Agar 100 resin. 70-nm sections were post-stained with 2% uranyl acetate and Reynolds lead citrate. Sections were examined with a FEI Morgagni 268D (FEI, Eindhoven, The Netherlands) operated at 80 kV. Images were acquired using an 11 megapixel Morada CCD camera (Olympus-SIS).

#### *In vivo* muscle permeability measurements

To determine in vivo muscle tissue permeability, we used the well described Evans blue dye assay ^47^Briefly, 1% Evans blue dye (10 ml kg^−1^ body weight, Sigma, GmbH) was injected into the intraperitoneal cavity of 8 month old animals. 16hours later mice were sacrificed via cervical dislocation and muscles collected. Quadriceps muscles were weighed and then soaked in formamide (GibcoBRL, UK) for 48 hr at 55°C with gentle shaking. The optical density of Evans blue in the resulting supernatant was measured at 610 nm with a Spectronic 610 spectrophotometer (Milton Roy). To image Evans blue via microscopy, muscles were immediately embedded in OCT and frozen. 10µm longitudinal sections were cut and visualized after counterstaining with DAPI.

#### Pcyt2 enzyme activity

Pcyt2 activity was measured as previously described ^17^. Briefly, frozen muscle tissue (50mg) was homogenized in a cold lysis buffer (10 mM Tris-HCl [pH 7.4], 1 mM EDTA, and 10 mM NaF) and briefly centrifuged to remove cell debris. Fifty ug of protein was assayed with 0.2 µCi of [14C]-phosphoethanolamine (P-Etn) (American Radiolabeled Chemical) in 50µl of reaction mixture of 50 mM MgCl2, 50 mM DTT, 10 mM unlabeled P-Etn, 20 mM CTP and 100 mM Tris-HCl (pH 7.8). The reaction was incubated at 37^0^C for 15 min and terminated by boiling (2 min). The reaction product [14C]CDP-Etn was separated from [14-C]PEtn on Silica gel G plates (Analtech) with a solvent system of methanol:0.5%NaCl:ammonia (10:10:1). CDP-Etn and P-Etn in standards and samples were visualized with a 0.5% ninhydrin in ethanol and the [14C]CDP-Etn collected and quantified by liquid-scintillation counting. Pcyt2 activity was expressed in nmol/min/mg protein. Protein content was determined with bicinchronic acid assay from Pierce.

#### Statistical analysis of mouse studies

All mouse data are expressed as mean +/-standard error of the mean (SEM). Statistical significance was tested by Student’s two tailed, unpaired t-test with Welch correction; Mann-Whitney U test; 1-way Multiple comparison ANOVA, with Welch correction, 2-way ANOVA followed by Bonferroni’s post-hoc test or Analysis of Co-Variance (ANCOVA) ^89^. All figures and mouse statistical analyses were generated using Prism 8 (GraphPad) or R. Details of the statistical tests used are stated in the figure legends. In all figures, statistical significance is represented as *\*P* <0.05, *\*\*P* <0.01, *\*\*\*P* <0.001, *\*\*\*\*P* <0.0001.

